# Improved Sperm Quality and Cryo-Induced Epigenetic Changes in Sterlet via Density-Gradient Sorting

**DOI:** 10.64898/2026.01.29.702533

**Authors:** Pavlina Vechtova, Anatolii Sotnikov, Jan Sterba, Borys Dzyuba

## Abstract

**Background:** Cryopreservation is a valuable tool in aquaculture and conservation programs, yet it exposes spermatozoa to physiological and molecular stresses that may impair motility, fertilisation capacity, and genomic stability. In fishes, where sperm motility is brief and easily activated, post-thaw separation of high-quality sperm remains technically challenging and poorly understood. This study evaluated whether density-gradient centrifugation can isolate a functionally superior subpopulation of sterlet (*Acipenser ruthenus*) spermatozoa with enhanced motility, fertilising ability, and reduced cryopreservation-induced epigenetic alterations. We further examined whether the use of this selected fraction influences DNA methylation patterns in resulting embryos.

**Results:** Cryopreservation substantially reduced the proportion of motile spermatozoa, while density-gradient centrifugation consistently enriched motile cells both before and after freezing. Motility enhancement reflected a higher proportion of cells capable of activation rather than changes in kinematic behaviour. Fertilisation trials confirmed that the selected post-thaw fraction produced fewer malformed embryos compared with unselected cryopreserved sperm. Cryopreservation and post-thaw selection induced small but significant methylation changes in sperm, predominantly in intergenic regions and promoter-proximal elements. However, these epigenetic differences were not maintained in embryos. Embryo methylomes showed minimal variation between treatments, no distinct clustering by sperm origin, and negligible numbers of differentially methylated regions. Thus, although cryopreservation and sperm selection influenced sperm DNA methylation, these alterations did not translate into measurable changes in embryo methylation patterns.

**Conclusions:** Density-gradient centrifugation effectively isolates a motile, functionally improved sterlet sperm fraction after cryopreservation, enhancing fertilisation outcomes and reducing developmental abnormalities. Cryopreservation and sperm selection introduce detectable but limited methylation variation in spermatozoa; however, these changes are not inherited by embryos. The findings highlight the utility of post-thaw sperm selection in aquaculture practice and indicate that cryopreservation-associated epigenetic variation in sterlet sperm does not propagate to early developmental stages.

## Introduction

Long-term storage of viable spermatozoa at low temperatures (cryopreservation or cryobanking) is a powerful tool having a high applicability potential in reproductive medicine, animal breeding, and conservation programs for preserving unique genetic resources and minimising the loss of biodiversity (Benson, et al., 2012; Watson, Holt, 2001; Yánez-Ortiz, et al., 2022). Despite the optimisation and widespread application of cryopreservation procedures across many animal groups, the technique still induces varying degrees of damage to certain sperm subpopulations, affecting their cell integrity, functionality, and genomic structure. To avoid the participation of cryo-damaged spermatozoa in fertilisation, post-thaw selection of non-damaged spermatozoa is commonly used. Even though these techniques are well-established for mammals, based on characteristics such as motility and a variety of physical properties (Pinto, et al., 2021), there is still room for future improvement (Ribas-Maynou, et al., 2022). Moreover, sperm selection tools are of great interest in cryobiology, as they enable direct assessment of cryopreservation-induced damage and help optimise freezing protocols by isolating viable sperm fractions for the next comprehensive analysis (Castro, et al., 2016). Notably, taxon-specific differences in sperm physical and physiological traits, shaped by distinct reproductive strategies, play a critical role in the design of both specific cryopreservation protocols and post-thaw selection methods for high-quality spermatozoa.

The wide diversity of reproductive modes and spawning environments among fishes results in considerable variability in their sperm physical, physiological and biochemical parameters, which predetermines the difficulties in sperm fractionation after cryopreservation. In addition, in all externally fertilising fishes, sperm motility is activated for a short time just after releasing sperm into the spawning environment before the gamete encounter (Dzyuba, Cosson, 2014). This short period of sperm motility necessitates avoiding motility activation during in vitro manipulation, including cryopreservation and post-thaw separation of spermatozoa. As a result, separating high-quality spermatozoa in fishes is a complex, often species-specific task, and the development of these methods is still in its early stages. That is why, nowadays, the massive knowledge of fish sperm cryopreservation and analysis of post-thaw genetic and biochemical alterations is an integral part of the entire sperm analysis. That is risky for the correct estimation of cryodamage, as only a few publications reported the application of cell separation techniques in spermatological studies in fishes (Bravo, et al., 2020; Li, et al., 2010; Valcarce, et al., 2016). The study by Horokhovatskyi et al. (Horokhovatskyi, et al., 2018) demonstrated that Percoll density-gradient centrifugation effectively isolates spermatozoa with high motility, viability, and minimal proteomic alterations after cryopreservation. However, the potential of this technique to minimise the risk of transmitting cryopreservation-induced mutations to the progeny has not yet been investigated, despite its clear relevance (Holt, 2023).

The assessment of risks associated with genetic alterations in progeny obtained using cryopreserved sperm in fishes has a relatively long history and is characterised by contradictory findings. To date, it is clear that sperm cryopreservation can have a positive (Bokor, et al., 2015; Panda, et al., 2024), negative (Dziewulska, et al., 2011; Nusbaumer, et al., 2019) or no (Dainat, et al., 2026; Linhart, et al., 2000; Miller, et al., 2018) detected effects on progeny growth parameters. These contrasting outcomes suggest strong species-specific responses that depend on the cryopreservation protocol applied, the developmental stage examined, and the criteria used for progeny evaluation (Bøe, et al., 2021; Depincé, et al., 2020). Considering that such effects may arise from sperm genetic damage (permanent DNA sequence alterations and fragmentation (Xin, et al., 2020)) as well as epigenetic modifications (heritable DNA modifications not involving changes in nucleotide sequence (Zhang, et al., 2023)), current understanding of the impact of sperm cryopreservation on genetic integrity and inheritance remains fragmented and requires further investigation. At the same time, DNA methylation is a known epigenetic mechanism, which evolved to facilitate rapid adaptation to immediate external stimuli of the host organism by regulation of appropriate cellular pathways on the level of gene expression (Feil, Fraga, 2012; Flores, et al., 2013; Law, Holland, 2019). That is why, considering cryopreservation as a significant source of stress, the study of cryopreservation-driven changes in sperm DNA methylation is relevant for the general understanding of sperm cryodamage.

Moreover, all related data on this topic in fish to date were obtained using the entire post-thaw sperm samples, without selecting “high-quality” spermatozoa, to obtain progeny that were minimally affected by sperm cryopreservation. Recent data on the possibility of selecting spermatozoa which are not only of significantly increased sperm motility and morphology, but also of decreased DNA genetic and epigenetic changes, performed in mammals (Ali, et al., 2022; Amano, et al., 2024; Lacalle, et al., 2023; Yu, et al., 2015) stimulate the broadening of this approach to fish, as nowadays it is not clear if this approach will provide the same positive result, especially taking into account essential differences in sperm structure and physiological properties between fish and mammals.

To address this knowledge gap, we hypothesised that density-gradient centrifugation represents an effective strategy for isolating a functionally superior subpopulation of cryopreserved sterlet spermatozoa characterised by enhanced motility, increased fertilisation capacity, and reduced cryopreservation-induced epigenetic alterations relative to the unselected sperm population. In this study, we examined whether cryopreservation-associated genetic and epigenetic damage is uniformly distributed across spermatozoa and whether targeted post-thaw selection of a high-motility fraction can improve sperm genetic integrity. Furthermore, we evaluated whether utilisation of this selected sperm fraction can mitigate cryopreservation-mediated declines in fertilising ability and reduce the incidence of embryonic malformations and epigenetic alterations in the resulting progeny.

## Materials and Methods

### Ethics

All maintenance procedures and handling of fish were conducted in accordance with approvals from the Institutional Animal Care and Use Committee at the University of South Bohemia. These approvals were based on authorisations for the breeding and supply of experimental animals (Reference numbers: 56665/2016-MZE-17214 and 64155/2020-MZE-18134) and on permission for the use of experimental animals (Reference number: 68763/2020-MZE-18134), issued to the Faculty of Fisheries and Protection of Waters by the Ministry of Agriculture of the Czech Republic.

### Fish maintenance and gametes collection

Sterlet (*Acipenser ruthenus*) broodstock (4–10-year-old, 45–65 cm standard length, 1.2–2.4 kg body weight) were maintained at the Genetic Fisheries Centre. Before experiments, breeders were moved from an outdoor pond (3 °C) to a recirculating system, where the temperature was raised to 16 °C at 1 °C/day. Gametes were obtained as described (Chebanov, Galich, 2011). Spermiation was induced by intramuscular injection of carp pituitary suspension (4 mg kg⁻¹) 32 h before sperm collection. Sperm was collected via catheter, stored on ice, and only samples with >80% motility were used. Females received two injections of carp pituitary: 0.5 mg kg⁻¹ (priming) and 4.5 mg kg⁻¹ (resolving) 12 h apart.

### General experimental design

The experimental design consisted of two experiments.

***Experiment 1*** was conducted to investigate the impact of sperm cryopreservation and separation of highly motile fraction using Percoll density gradient centrifugation (DGC) on sperm motility parameters and DNA methylation state (Fig. 1). Sperm from 10 males were used to assess motility parameters. Sperm samples from three of those males were also used to evaluate DNA methylation using whole-genome bisulfite sequencing (WGBS).

**Fig. 1.**
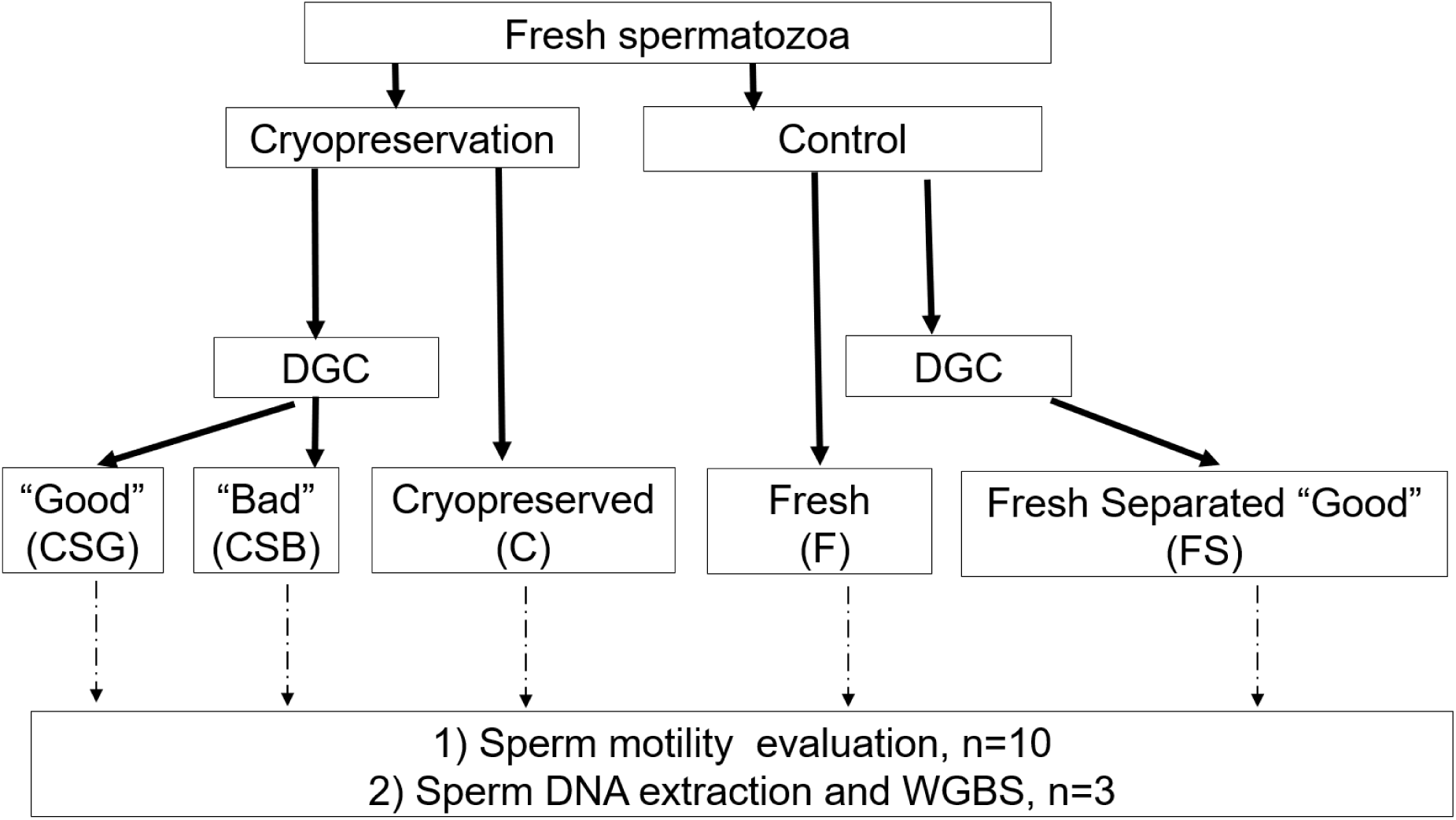
Experimental design (Experiment 1) for sperm sample processing and analysis. Fresh spermatozoa (from 3 males) were allocated to two experimental conditions: cryopreserved and fresh control. Cryopreserved samples were analysed directly (C) or separated by density gradient centrifugation (DGC) into “Good” (CSG) and “Bad” (CSB) fractions. Fresh samples were analysed without processing (F) or after DGC to obtain the “Good” fraction (FS). In samples F, FS, C, and CSG, sperm motility assessment by CASA was performed in samples from 10 males (n=10) and DNA methylation analysis by whole-genome bisulfite sequencing (WGBS) was performed in all samples from 3 males (n=3).

Fresh spermatozoa from 3 males each were divided into two experimental subgroups – before and after cryopreservation. After cryopreservation, samples were either analysed directly as cryopreserved (C), representing the unprocessed post-thaw state of whole ejaculate, or subjected to density gradient centrifugation (DGC) to separate sperm fractions. This separation yielded two fractions: “Good” (CSG), enriched in motile sperm, and “Bad” (CSB), containing mainly immotile sperm. This step enables evaluation of whether post-thaw DGC allows selection of a sperm fraction with minimal changes in control sperm quality and epigenetic integrity.

Fresh sperm (F) was also analysed directly without cryopreservation to serve as a baseline for motility and DNA methylation. Additionally, fresh samples underwent DGC to obtain the “Good” fraction (FS), enabling comparison of selection effects in fresh versus cryopreserved conditions.

The experimental groups (C, CSG, F, and FS) were subjected to sperm motility evaluation by CASA and further CASA data examination. All five experimental groups (C, CSG, CSB, F, and FS) were subjected to sperm DNA extraction followed by WGBS to investigate epigenetic modifications associated with cryopreservation and sperm selection.

***Experiment 2*** was designed to investigate the sperm fertilising ability and DNA methylation state in embryos (Fig. 2) produced using sperm samples that underwent different treatments described in Experiment 1 (cryopreservation and selection of sperm with high motility parameters).

**Fig. 2.**
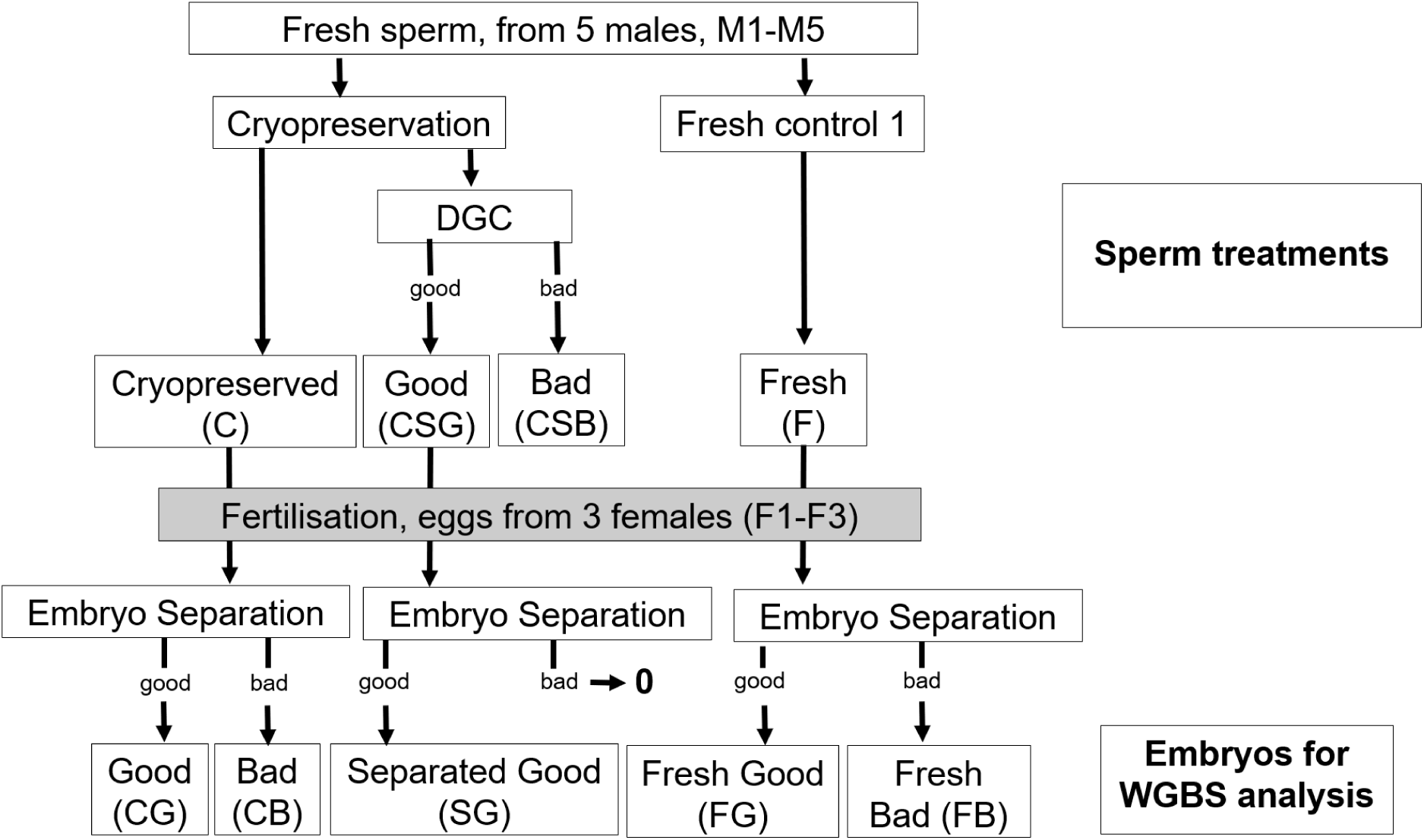
Experimental design for embryo DNA methylation analysis (Experiment 2). Fresh spermatozoa from 5 males (M1–M5) were processed using the same treatments as in Experiment 1. For fertilisation, the only fresh control (F), cryopreserved (C), and cryopreserved and separated by DGC “good” fraction (CSG) were used. These sperm groups were used to fertilise oocytes from 3 females (F1–F3). Resulting embryos were classified into “Good” or “Bad” groups based on external features: CG and CB (from C), SG (from CSG), FG and FB (from F). All embryo groups were analysed for DNA methylation using whole-genome bisulfite sequencing (WGBS).

The following sperm samples from 5 males (M1–M5) were used to obtain progeny: fresh, unprocessed sperm (F), Cryopreserved (C), cryopreserved and DGC separated (CSG). These sperm groups were used to fertilise individual batches of oocytes from three females (F1–F3). After hatching, the embryos from each experimental group were further classified as normally developing (“Good”) and malformed (“Bad”) based on visual morphometric appearance. Hereby, the groups of embryos for WGBS were: “Good” embryos obtained with whole cryopreserved sperm - CG, “Bad” embryos obtained with whole cryopreserved sperm -CB, good embryos obtained with a fraction of cryopreserved sperm after DGC – SG, and Good and Bad embryos obtained using fresh sperm FG and FB, respectively.

All these embryo groups (CG, CB, SG, FG, FB) were subjected to DNA methylation analysis using Whole-Genome Bisulfite Sequencing (WGBS).

### Sperm motility analysis

Sperm motility was activated in a medium containing 1 mM CaCl₂, 10 mM Tris-HCl (pH 8.0), and 0.25% Pluronic F-127 (P2443, Sigma-Aldrich/Merck) to prevent adhesion to the glass slide. Motility was recorded in the ISAS spermtrack-10 sperm counting chamber (PROISER, Spain) for 120 s after motility activation, using a negative phase-contrast microscope with x10 objective (LM 666 PC, PROISER, Spain), and an IDS digital camera (UI-3130CP-M-GL, Imaging Development Systems GmbH, Germany), at 50 fps frame rate. The videos were analysed using the CASA automated plugin for ImageJ v.1.54g (Purchase, Earle, 2012; Wilson-Leedy, Ingermann, 2007) to determine: motile cell percentage (MOT, %), VCL (curvilinear velocity, µm s⁻¹), VAP (average path velocity, µm s⁻¹), VSL (straight-line velocity, µm s⁻¹), LIN (linearity, VSL/VAP), WOB (track oscillation, VAP/VCL), and BCF (beat-cross frequency, Hz). Statistical analysis was performed starting at 15 s post-activation, when sample drift was no longer observed.

### Sperm cryopreservation and thawing

Sperm was diluted 1:1 in an extender containing 30 mM Tris-HCl, 23.4 mM sucrose, and 0.25 mM KCl with 15% methanol (modified after (Horváth, et al., 2008a). Diluted samples were packed in 0.5 mL plastic straws, incubated on ice for 10 min, then placed on a 3 cm polyester raft in a Styrofoam box over liquid nitrogen for 10 min. Straws were then plunged into liquid nitrogen for storage. Thawing was performed in a 40 °C water bath for 6 s (Dzyuba, et al., 2010).

### Sperm fractionation by Percoll density gradient centrifugation

Percoll density gradient centrifugation (Percoll DGC) was performed as described (Horokhovatskyi, et al., 2018). Briefly, a 15 mL tube was prepared with two 2 mL Percoll layers (90% and 40%), and 1 mL of sperm was layered on top. Percoll solutions were prepared in a buffer containing 16 mM NaCl, 3 mM KCl, 0.19 mM CaCl₂, and 10 mM Tris-HCl (pH 8.0), corresponding to non-activating medium (NAM) and native sterlet seminal fluid. Tubes were centrifuged at 2,000 × g for 10 min at 4 °C. After centrifugation, the upper fraction consisted mainly of immotile spermatozoa, whereas the bottom 0.5 mL fraction contained predominantly motile spermatozoa. The collected fractions were transferred to 2 mL tubes, diluted with 1.5 mL NAM, and centrifuged at 1,000 × g for 10 min at 4 °C. Pellets were resuspended in 0.5 mL NAM. Following Percoll DGC of fresh sperm samples, the upper fraction contained insufficient sperm numbers and was therefore excluded from further analyses. The bottom fraction of fresh sperm, consisting of motile spermatozoa, was designated FS. After cryopreservation and Percoll DGC, both upper and bottom fractions contained sufficient sperm for downstream analyses and were designated CSB and CSG, respectively.

### Fertilisation trials

To assess fertilisation outcomes using cryopreserved sperm before and after separation, experimental and control sperm samples were mixed with ∼150 eggs (≈2 g) and 8 mL hatchery water for 2 min. Sperm volume was adjusted to achieve an egg-to-sperm ratio of 1:50,000. Sperm concentration was determined using a Bürker chamber after 1:2000 dilution in physiological solution. Fertilised eggs were incubated in Petri dishes in a flow-through incubator operated in recirculation mode at 15 °C for 7 days. At 72 h post-fertilisation, embryo development was assessed from photographs. After hatching, larvae were anaesthetised with 100 mg L⁻¹ MS-222 (E10521, Sigma-Aldrich/Merck) to evaluate hatching and embryo malformation rates as percentages of fertilised eggs. The collected data were used for graphical presentation and statistical analysis.

### Sperm DNA extraction and whole genome bisulfite sequencing (WGBS)

#### DNA extraction from sperm

Genomic DNA samples were isolated from sperm using the NucleoSpin Tissue kit (Macherey-Nagel) according to the modified NucleoSpin Tissue protocol for semen. A 25 µL sperm sample (for fresh sperm) or a 50 µL sample (for all other types) was diluted with 400 µL of pre-warmed B3 and T1 buffers, in equal volumes. After vertexing, 50 µL of Proteinase K was added, the samples were vortexed again, and the mixture was incubated for 10 minutes at 70 °C. After the initial incubation, the samples were vortexed again and incubated overnight at 60 °C. The rest of the procedure was done according to the manufacturer’s recommendation. Changes for the first step were made because the original protocol wasn’t suitable for extracting DNA from sturgeon spermatozoa.

#### DNA extraction from embryo

Genomic DNA samples were isolated from embryo using the NucleoSpin Tissue kit (Macherey-Nagel) according to manufacturers instructions. The tissue lysis step was prolonged to 3 hours, the elution step was repeated twice with 2x 50 μl of elution buffer.

#### Bisulfite conversion, sequencing library preparation and sequencing

Sperm gDNA was bisulfite-converted, and a sequencing library was prepared using Zymo EZ DNA Methylation Gold Kit and Swift Accel-NGS MethylSeq kit for directional sequencing by Admera Health Biopharma Services. The sequencing was done on NovaSeq X Plus 2x150 (Illumina, San Diego, CA, USA).

*Embryos* gDNA was bisulfite-converted, and a sequencing library was prepared using EpiTect Fast bisulfite conversion kit and QIAseq Methyl Library Kit for directional sequencing by UCLA Technology Center for Genomics & Bioinformatics (TCGB). The sequencing was done in the NovaSeq X Plus 25B sequencer at PE 150 cycles (Illumina, San Diego, CA, USA).

### Data analysis

#### Sperm motility parameters

Obtained after CASA dataset 1 contained: 1) data on motility percentage evaluated at each second post-activation (15-120s post-activation, 105 time-points) and associated with male ID (n=10), experimental condition (n=4) and 2) the kinematic parameters obtained from all tracked spermatozoa (n=1.2x10^6^) associated with the above indices. After z-score standardisation, the data set was subjected to cluster analysis. The optimal cluster number was determined by k-means clustering (k = 2–10) using average silhouette width on random subsamples, repeated ten times with different seeds. The most frequent k was applied to the full dataset for final clustering. Cluster characteristics (mean ± SE) were summarised, and relative cluster abundance per treatment and male was calculated and further analysed by two-way ANOVA.

The dataset 1 was also used to create the next dataset (Dataset 2), containing averaged motility parameters for each male (n=10) at each experimental condition and each second post-activation. Dataset 2 was used for the general presentation of the dynamics of each sperm motility parameter (15-120s post-activation, 105 time points), and for the two-way repeated-measures ANOVA at each post-activation time point.

Two-way repeated-measures ANOVA (after confirming normality via Kolmogorov–Smirnov test, p ≥ 0.05) with Bonferroni post-hoc tests (p < 0.05 considered significant) to assess differences in motility parameters. Factors: cryopreservation (CRYO: control (F) vs. cryopreserved (C) groups) and fractionation (DGC: before vs. after DGC treatment), forming four groups: F, FS, C, and CSG (see Fig. 1). Significant interactions were followed by simple-effects comparisons; otherwise, main effects were interpreted.

#### Embryo development analysis

A nonparametric repeated-measures analysis using the Friedman ANOVA, followed by the Wilcoxon matched-pairs test with Bonferroni correction, was used to evaluate embryo hatching and abnormality rates.

#### WGBS data analysis

##### Raw data processing, mapping and BS-seq analysis

Raw read quality was evaluated using fast QC (Andrews, 2010). Raw reads were trimmed off adapters and filtered for low-quality reads using Trimmomatic v0.38 (Bolger, et al., 2014) with the following parameters: ILLUMINACLIP seed mismatches = 2, palindrome clip threshold = 30, simpleClipThreshold = 10; LEADING and TRAILING = 3, MINLEN = 36. The clean reads were mapped to the *A. ruthenus* reference genome (GCF_902713425.1) with Bismark v0.24.0 (Krueger, Andrews, 2011) in PE mode with a relaxed mapping stringency --score_min L,0,-0.6 and collecting unmapped and ambiguously mapped reads to separate files with --un and –ambiguous options. Other parameters were default. Unmapped reads were remapped in SE mode. Methylated cytosines in all contexts were extracted from PE and SE bam files with Bismark’s bismark2bedGraph script. Genome coverage for each sample was calculated and plotted using the plotCoverage tool from deepTools v3.3.0 (Ramírez, et al., 2016), and the deepTools coverage summary was visualised using ggplot2 (Wickham, 2016). Pairwise differential methylation was calculated with Defiant v1.1.9 (Condon, et al., 2018). The Defiant run was modified using the options -CpN 5, -c 10, -d 4, -P 10, -v bh, -E; other parameters were set to their default values.

##### Genome annotation

Missing genomic features in the reference genome annotation file were inferred using several tools. Introns, intergenic regions, start and stop codons were inferred using agat v0.8.0. and v1.4.1 (Dainat, et al., 2026). Promoters were identified with the ‘promoters’ function of the GenomicRanges package (Lawrence, et al., 2013). Promoter coordinates were defined as 1500 bp upstream and 500 downstream of TSS. UTRs were derived using UCSC Genome Browser’s utilities gtfToGenePred and genePredToBed (Kent, et al., 2002), followed by NCBI’s Add_UTRs_to_GFF.py public script (retrieved from ftp://ftp.ncbi.nlm.nih.gov/genomes/TOOLS/add_utrs_to_gff/). CpG islands and shores were identified using Emboss tools (Rice, et al., 2000). CpG shores were defined as regions 2000 bp upstream and downstream of the CpG islands (Irizarry, et al., 2009; Sandoval, et al., 2011).

##### Differentially methylated region annotation

Differentially methylated regions (DMRs) of each pairwise comparison were mapped and quantified over annotated genomic features using the GenomicRanges and Genomation packages (Akalin, et al., 2014). Results were visualised using the ggplot2 package. DMG intersection among all comparisons was assessed in R. The Upset plot for multiple-group intersections visualisation was constructed using the UpsetR package (Conway, et al., 2017).

##### Principal component analysis

Data integration for principal component analysis (PCA) was performed with methylKit v1.36.0 (Akalin, et al., 2012). The CpG counts extracted with Bismark’s bismark2bedGraph script were processed with the filterByCoverage function, removing bases represented by fewer than 10 reads, which are considered unreliable, and bases above the 99.9th percentile of coverage in each sample, probably representing PCR bias or overamplification. Coverage was normalised to adjust for differences in coverage across samples, using a scaling factor derived from the difference in the median coverage distribution between samples, as implemented in the normalizeCoverage function. Results were visualised using the scatterplot3d function of the plotly package v2.11.1 (Li, Bilal, 2021).

##### Gene set enrichment analysis (GSEA)

The GSEA was done using Goatools. The GAF file derived from the *A. ruthenus* reference genome and the *A. ruthenus* background gene set were downloaded from NCBI ftp on 11 Dec 2024. The obo file, data version 2024-10-27, was downloaded on 12 Nov 2024.

## Results

### Sperm motility prior to and after cryopreservation and fractionation

#### Sperm motility percentage

Across the post-activation period, MOT showed the clearest and most consistent treatment effects among all analysed sperm motility variables. The temporal profile of average MOT for each experimental group and summary of statistical analysis are shown in Fig. 3. During the early and mid-motility interval (15–98 s, interval A, Fig 3), CRYO and DGC main effects and CRYO×DGC interaction were significant at every time point.

**Fig. 3.**
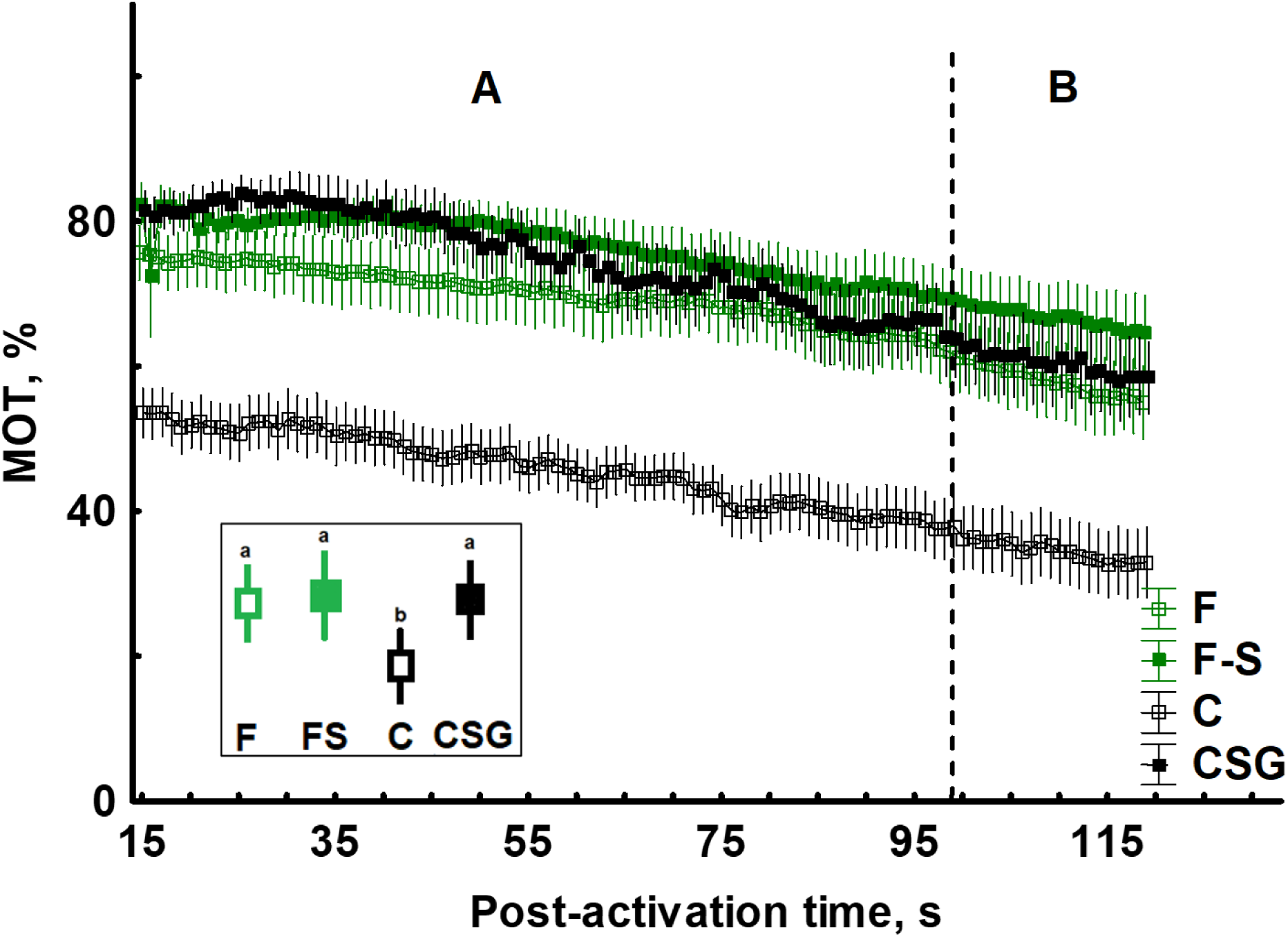
Post-activation dynamics of sperm motility. Values are presented as mean ± SE (n=10) for the four experimental groups: F (before cryopreservation and separation by Percoll), FS (before cryopreservation, after separation by Percoll), C (after cryopreservation and no separation by Percoll) and CSG (after cryopreservation and separation by Percoll). The dashed line indicates two time periods at which: A – the main effects of DGC, CRYO, and CRYO×DGC interaction are significant, with exceptions at time points marked by white rectangles at time axis, insertion indicates main trends of between groups comparison, where values with different letters differ significantly (Bonferroni p < 0.05); B – the only main effects of PERCOL and CRYO are significant.

The motility percentage in the samples after cryopreservation (C) was significantly lower than that of other sperm samples. The treatment of cryopreserved sperm with Percoll centrifugation (CSG) increased the MOT percentage to a level similar to that of the F and FS samples.

From 99 s onwards (Fig. 3), the CRYO×DGC interaction became consistently non-significant, with only isolated exceptions at a few individual time points. In this late motility interval (99–120 s), post-hoc tests therefore followed the interpretation of main effects only. At all these time points, both the CRYO and DGC main effects remained significant. Pairwise comparisons of estimated marginal means showed a persistent decrease in MOT in cryopreserved samples relative to fresh samples, and a consistent improvement in motility following Percoll treatment, independent of cryopreservation.

#### Kinematic parameters (VCL, VAP, VSL, LIN, WOB, BCF)

In contrast to MOT, the six kinematic parameters exhibited only sporadic significant effects of the experimental factors. The velocity-related parameters (VCL, VAP, and VSL) showed significant effects of CRYO, DGC, or their interaction at only a few, isolated time points, without forming clear temporal patterns. This is exemplified in VAP (Fig. 4a), where the maximum number of these time points was detected at the end of motility observation. Because of the rare effects of treatments and their interactions during the motility period, no further post-hoc analysis was performed for VCL, VSL, and VAP. For WOB, a consistent and significant interaction of factors was found at the later stage of the motility period only (Fig. 4b). For LIN and BCF, significant effects of CRYO, DGC, or their interaction were observed only at isolated time points, without forming a clear temporal pattern; therefore, further post-hoc comparisons were not performed for these parameters.

**Fig. 4.**
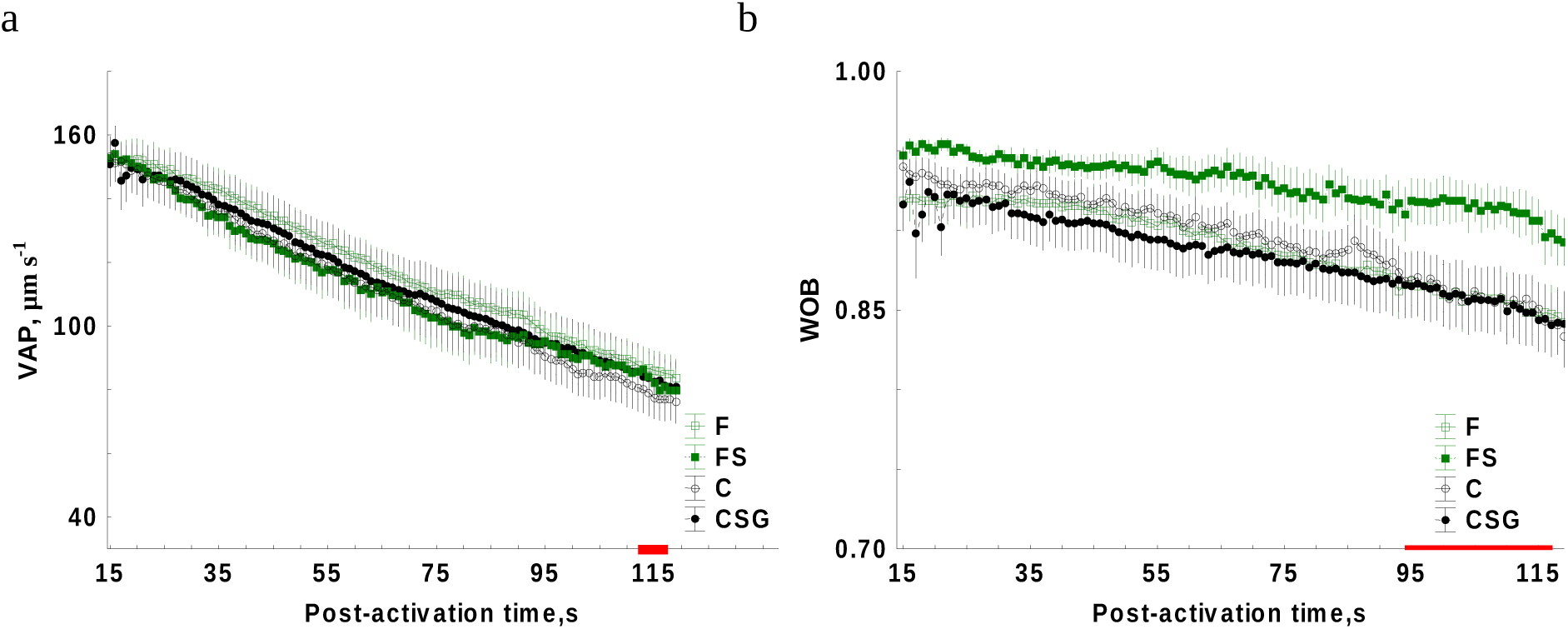
Examples of post-activation dynamics of sperm motility parameters in experimental groups. Values are presented as mean ± SE (n = 10) for the four experimental groups: F – before cryopreservation and separation by Percoll, FS – before cryopreservation, after separation by Percoll, C – after cryopreservation and no separation by Percoll and CSG – after cryopreservation and separation by Percoll. (a) – average path velocity (VAP), significant effects of the factors were detected only at isolated individual time points marked by red rectangles; (b) – oscillation of the track (WOB), significant effects of interaction of factors were detected in the post-activation time period marked by red colour.

Therefore, none of these parameters demonstrated treatment effects comparable in magnitude or consistency to those observed for MOT. Overall, the kinematic characteristics of motile spermatozoa remained largely stable across treatments.

#### Sperm cluster dynamics

Cluster analysis, performed based on the combined set of kinematic parameters (VCL, VAP, VSL, LIN, WOB and BCF), allowed to describe sperm heterogeneity by assigning motile spermatozoa to four motility classes (Fig. 5): Cluster 1 – spermatozoa with average velocities and low linearity; Cluster 2 – spermatozoa with average velocities and high linearity; Cluster 3 – spermatozoa with high velocities and high linearity; and Cluster 4 – spermatozoa with low velocities and high linearity.

**Fig. 5.**
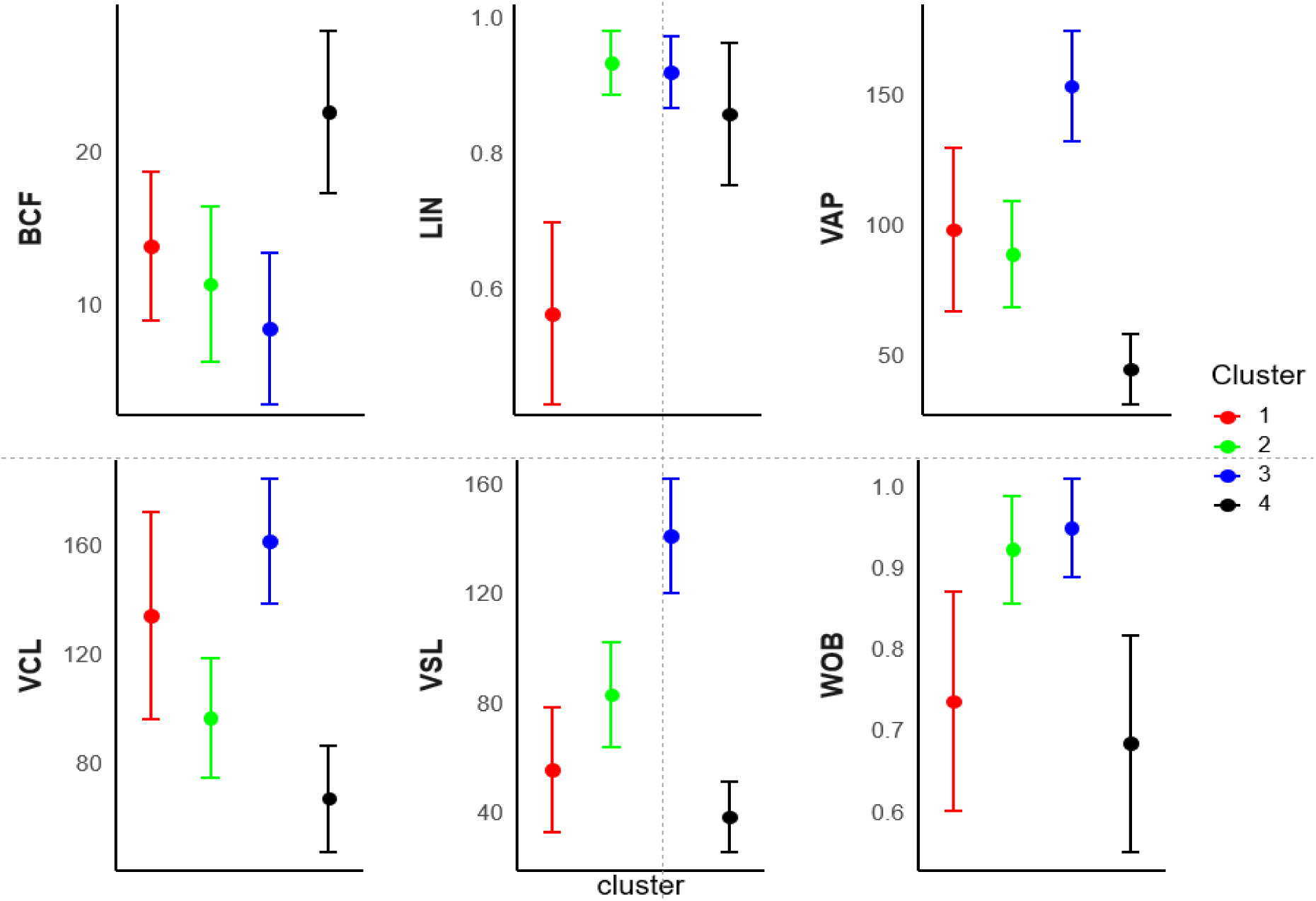
Description of kinematic characteristics for the sperm clusters determined by CASA analysis of all experimental samples. Data are presented after analysis of ∼ 1.2x10^6^ spermatozoon tracks from all experimental conditions and post-activation times, as mean ± SE.

A very similar general tendency in cluster dynamics was observed across experimental groups. In all treatments, the abundance of Cluster 3 decreased progressively over post-activation time, accompanied by a compensatory increase in the abundances of Clusters 2 and 4, while the abundance of Cluster 1 remained very low and did not change during the motility period. Cluster-abundance dynamics for the C and CSG groups are shown as examples in Fig. 6a and Fig. 6b, respectively.

**Fig. 6.**
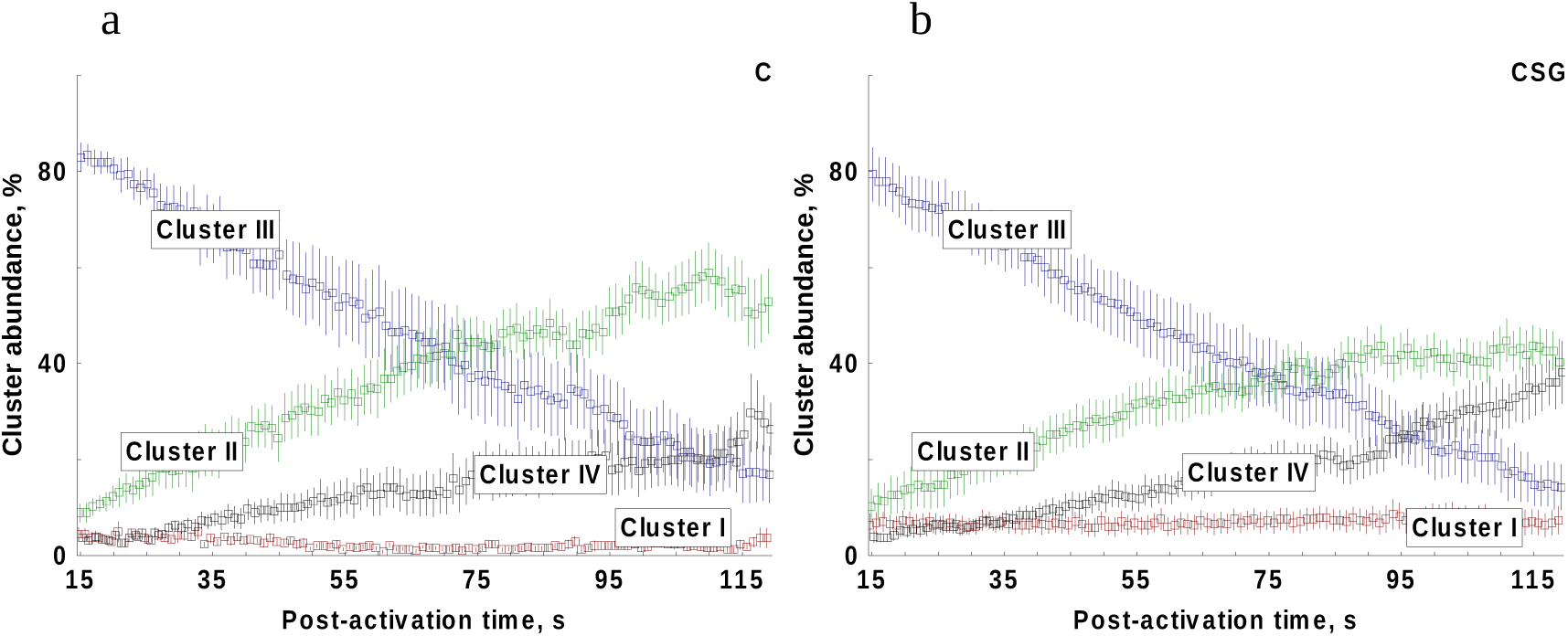
Examples of outcomes from cluster analysis of sperm motility parameters. Values are presented as mean ± SE (n = 10). (a) – cluster abundance in samples experimental group C (sperm after cryopreservation); (b) – cluster abundance in samples CSG (after cryopreservation and Percoll DGC treatment).

A two-way repeated-measures ANOVA of each cluster abundance at each post-activation time point revealed that significant effects of CRYO, PERCOLL and/or the CRYO×PERCOLL interaction were only sporadically detected at a few isolated time points. As there was no evidence of systematic treatment effects on cluster abundances, no further post-hoc tests were conducted.

Altogether, MOT was the parameter most strongly influenced by cryopreservation and Percoll treatment, whereas the kinematic characteristics of motile spermatozoa showed minimal and inconsistent effects of treatment. Cryopreservation consistently reduced the proportion of motile spermatozoa, whereas Percoll increased it by selectively eliminating cells unable to activate motility. Cluster analysis supported this interpretation, indicating that Percoll primarily improves sample quality by enriching the motile fraction rather than modifying the intrinsic swimming behaviour of motile spermatozoa.

### Embryo hatching and abnormality rates

Performed analysis of hatching rates obtained during the experiments, found no significant differences among experimental groups (Fig. 7a, Friedman ANOVA, Chi Sqr. = 2.27, p = 0.32). Based on the visual detection of abnormalities in the external morphology of hatched larvae (Fig. 7c), a significantly lower abnormality rate was found in larvae obtained with CSG sperm in comparison to C and F groups, while no difference was found between larvae obtained with sperm from experimental groups F and C (Fig. 7b).

**Fig 7.**
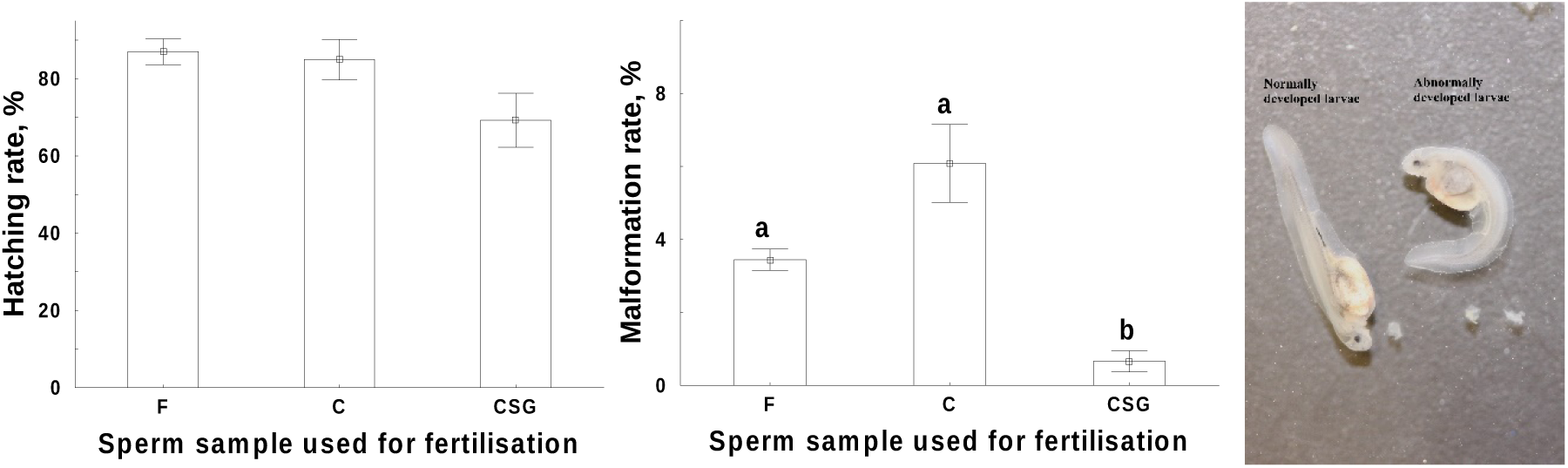
The outcomes of experimental sperm samples application in fertilisation tests. (a) – Hatching rate in progeny obtained with experimental sperm samples. No significant differences among groups were found (Friedman ANOVA, p = 0.32); (b) Malformation rate of abnormally developed larvae. The data are presented as mean ± SD. Values with different capital letters are significantly different (Wilcoxon matched-pairs test with Bonferroni correction, p < 0.05). Abbreviations for sperm samples used for fertilisation: F – before cryopreservation and separation by Percoll, C – after cryopreservation and no separation by Percoll and CSG – after cryopreservation and separation by Percoll.

### Whole genome bisulfite sequencing (WGBS)

#### Experiment 1

##### Bisulfite sequencing of sterlet spermatozoa DNA

Five sterlet spermatozoa experimental groups, Fresh (F), Fresh Separated (FS), Cryopreserved (C), Cryopreserved Separated Good (CSG) and Cryopreserved Separated Bad (CSB) were submitted to WGBS, each represented by three biological replicates. The experimental design is outlined in Fig. 1. We received, on average, 175M raw and 159M clean reads for sperm samples, with an average mapping efficiency of 73.5%. Details of raw sequencing data quality, raw-read adapter-trimming and quality-filtering, clean read mapping and deduplication statistics are given in Supplementary Tables S1–S3. The all-sample mean read coverage after deduplication is 8.8 x. The graphical representation of the read coverage distribution for each sample is available in Supplementary Fig. S1. The frequency of read coverage distribution for each sample is presented in Supplementary Fig. S2.

The average percentage of methylated cytosines in CpG context (mCpG) in all experimental groups is 82% (Fig. 8), which aligns with typical vertebrate genomes (Elango, Yi, 2008; Tweedie, et al., 1997). An overview of cytosine methylation in all dinucleotide contexts is listed in Supplementary Table S4.

**Fig. 8.**
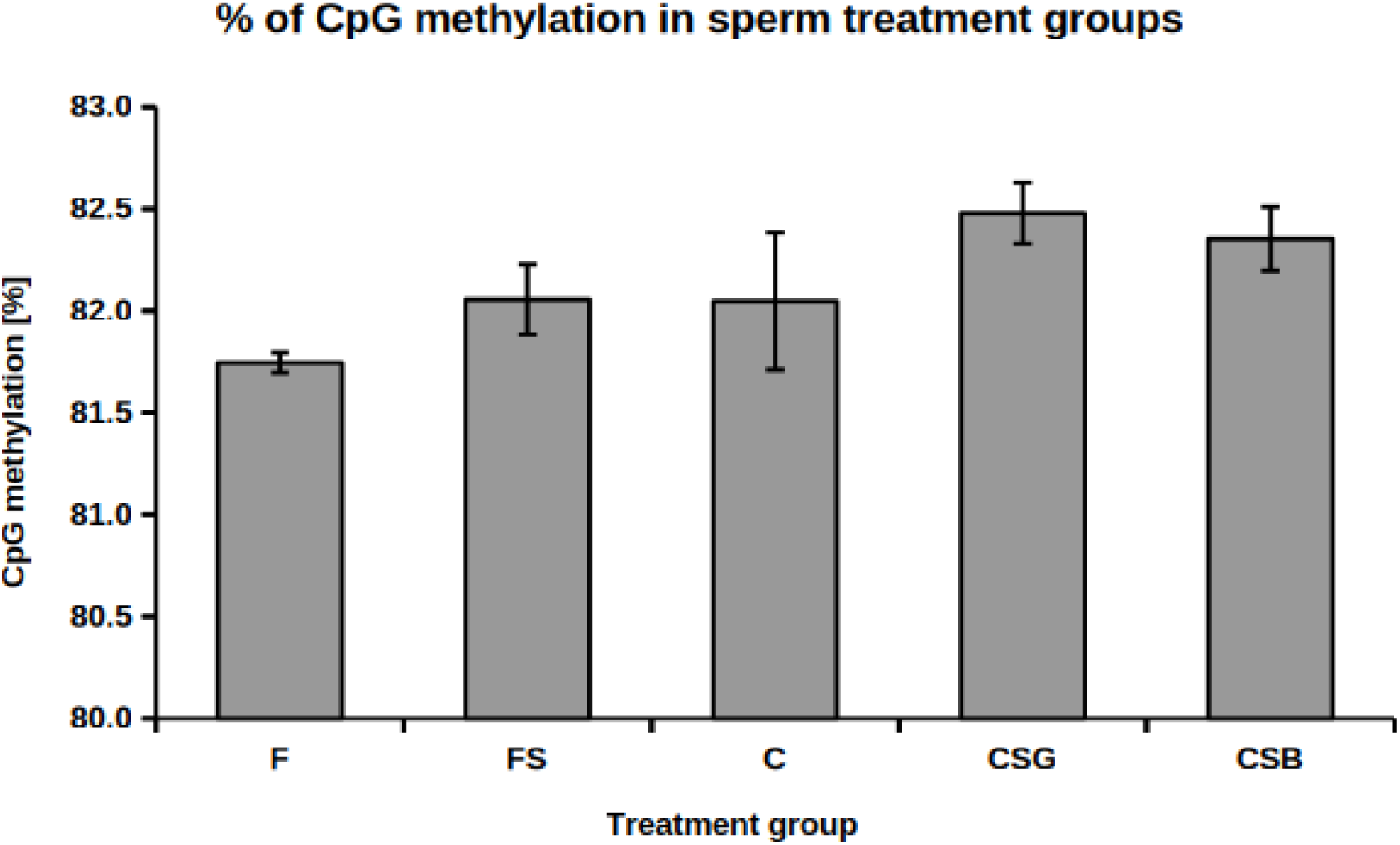
Percentage of CpG methylation in each treatment group, as summarised by the Bismark mapping report. Each value is represented by three biological replications. Treatment groups: F – Fresh, FS – Fresh Separated, C – Cryopreserved, CSG – Cryopreserved Separated Good, CSB – Cryopreserved Separated Bad.

The variance in CpG methylomes among experimental groups was further investigated in relation to cryopreservation and separation of different physiological forms. A principal component analysis of the median-normalised sample matrix of mCpG counts was projected in three dimensions (Fig. 9).

**Fig. 9.**
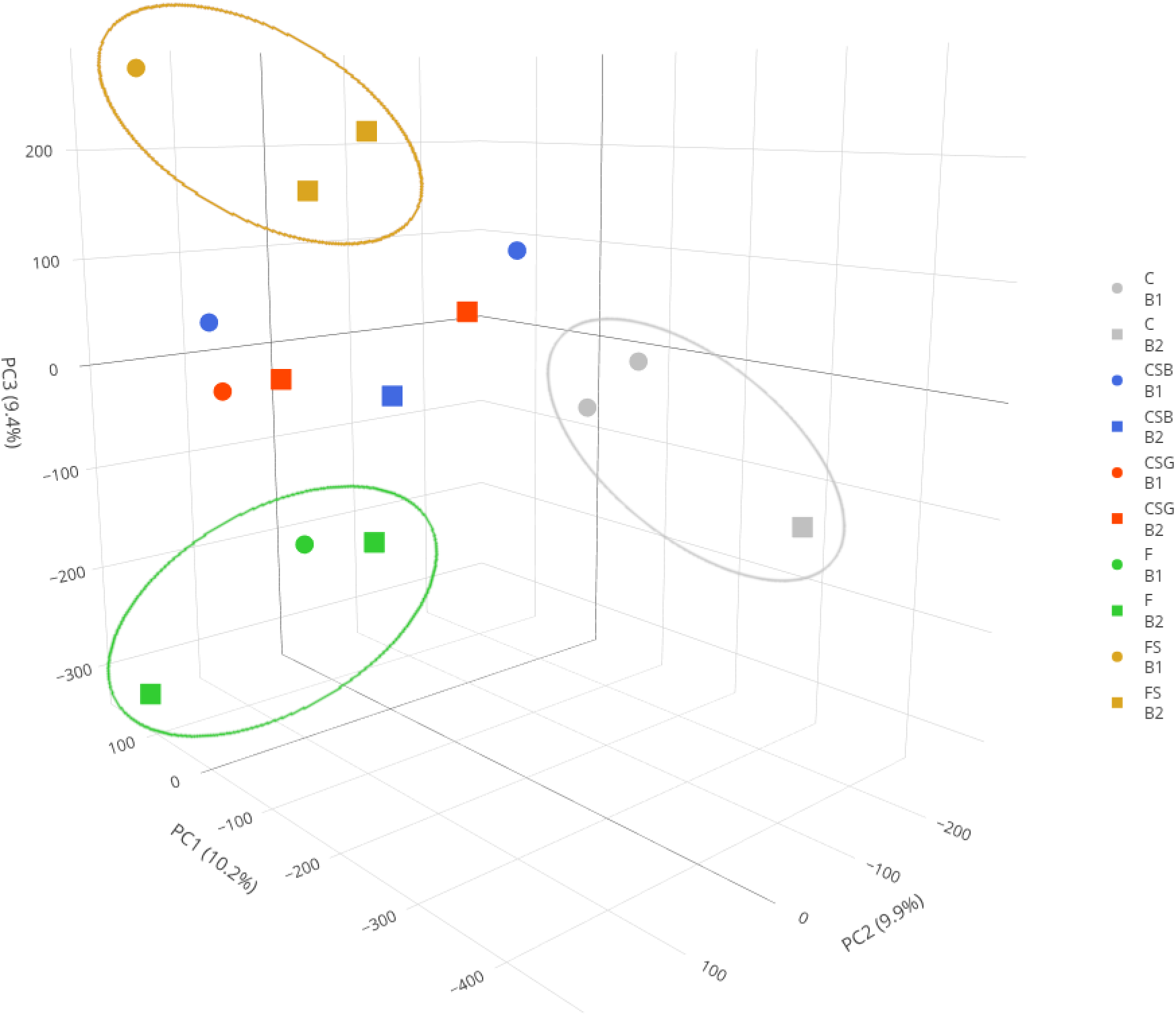
PCA of mCpG counts plotted along 3 major principal components in a 3D scatter plot. The mCpG count matrix of the spermatozoa experimental group was between-sample median-normalised before PCA calculation and plotting. Data points are sample-wise colour-coded and batch effect affiliation is discriminated by data point shape; circle – B1 batch group, rectangle – B2 batch group. Axis labels show % of data variance explained by each PC dimension.

Normalised mCpG counts plotted in the 3D space of principal components revealed the major source of methylome variance among experimental groups. 3D PCA enabled discrimination of 3 distinct clusters representing biological replicates of C, F, and FS groups. CSB and CSG groups data points were randomly distributed in the 3D PCA data space. mCpG count variance introduced by the potential batch effect was discriminated by data point shapes.

##### Identification and quantification of differentially methylated regions

Differentially methylated regions (DMR) in the CpG context were identified in a pair-wise comparison of 10 spermatozoa experimental group pairs. The methylation per cent change and corresponding p-value for each comparison, along with the total number of DMRs and their association with genes and intergenic regions, are presented in Table 1.

**Table 1.** Per cent change of differentially methylated regions (DMRs) for each pairwise comparison as calculated by Defiant*.

| control group | treatment group | percent change [%] | p-value | significance level | tot DMRs | DMRs genes | DMRs intergenic |
| --- | --- | --- | --- | --- | --- | --- | --- |
| F | FS | 0.27 | 0.0147 | * | 83 | 25 | 60 |
| F | C | 0.30 | 0.1265 |  | 78 | 29 | 52 |
| F | CSG | 0.72 | 0.0077 | ** | 18 | 6 | 11 |
| F | CSB | 0.60 | 0.0132 | * | 18 | 8 | 9 |
| FS | C | 0.03 | 0.8356 |  | 79 | 30 | 49 |
| FS | CSG | 0.45 | 0.0094 | ** | 23 | 3 | 19 |
| FS | CSB | 0.33 | 0.0278 | * | 18 | 6 | 9 |
| C | CSG | 0.42 | 0.0488 | * | 21 | 4 | 16 |
| C | CSB | 0.30 | 0.1060 |  | 13 | 7 | 7 |
| CSG | CSB | -0.12 | 0.3161 |  | 6 | 2 | 4 |

\* Each comparison, represented by three biological replications, is supported by a corresponding p-value and significance level (\* $p < 0.05$ , \*\* $p < 0.01$ ). Absolute total DMR numbers and separate DMR numbers associated with genes and intergenic regions are also listed. Treatment groups: F – Fresh, FS – Fresh Separated, C – Cryopreserved, CSG – Cryopreserved Separated Good, CSB – Cryopreserved Separated Bad.

The pairwise comparison of CpG methylomes between these experimental groups was performed to capture methylation changes possibly induced by cryopreservation and/or associated with different physiological forms in both cryopreserved and fresh spermatozoa populations. mCpG per cent change in any of the comparisons did not exceed 1 %, and besides the CSGxCSB comparison was always positive. The methylome changes in FxC, FSxC, CxCSB and CSGxCSB comparisons were not significant.

##### Annotation of DMRs

DMRs from all pairwise comparisons were mapped to the most relevant known genomic features (genes, exons, lncRNA) and to newly identified genomic features in this study (introns, 3’UTRs, 5’UTRs, promoters, intergenic regions, CpG islands, CpG shores). Absolute counts of DMRs for each genomic feature, as well as their relative ratios within each comparison, were calculated using the Genomation package and plotted in the form of a clustered heat map showing both relative and absolute DMR counts for each comparison and genomic feature (Fig. 10). Relative DMR ratios include DMRs, which are shared by overlapping genomic features and thus the row-wise ratios sum up above 100% per each comparison. The distribution of absolute mCpG counts in genomic features for all comparisons was visualised in a multi-facet pie chart in Supplementary Fig. S3.

**Fig. 10.**
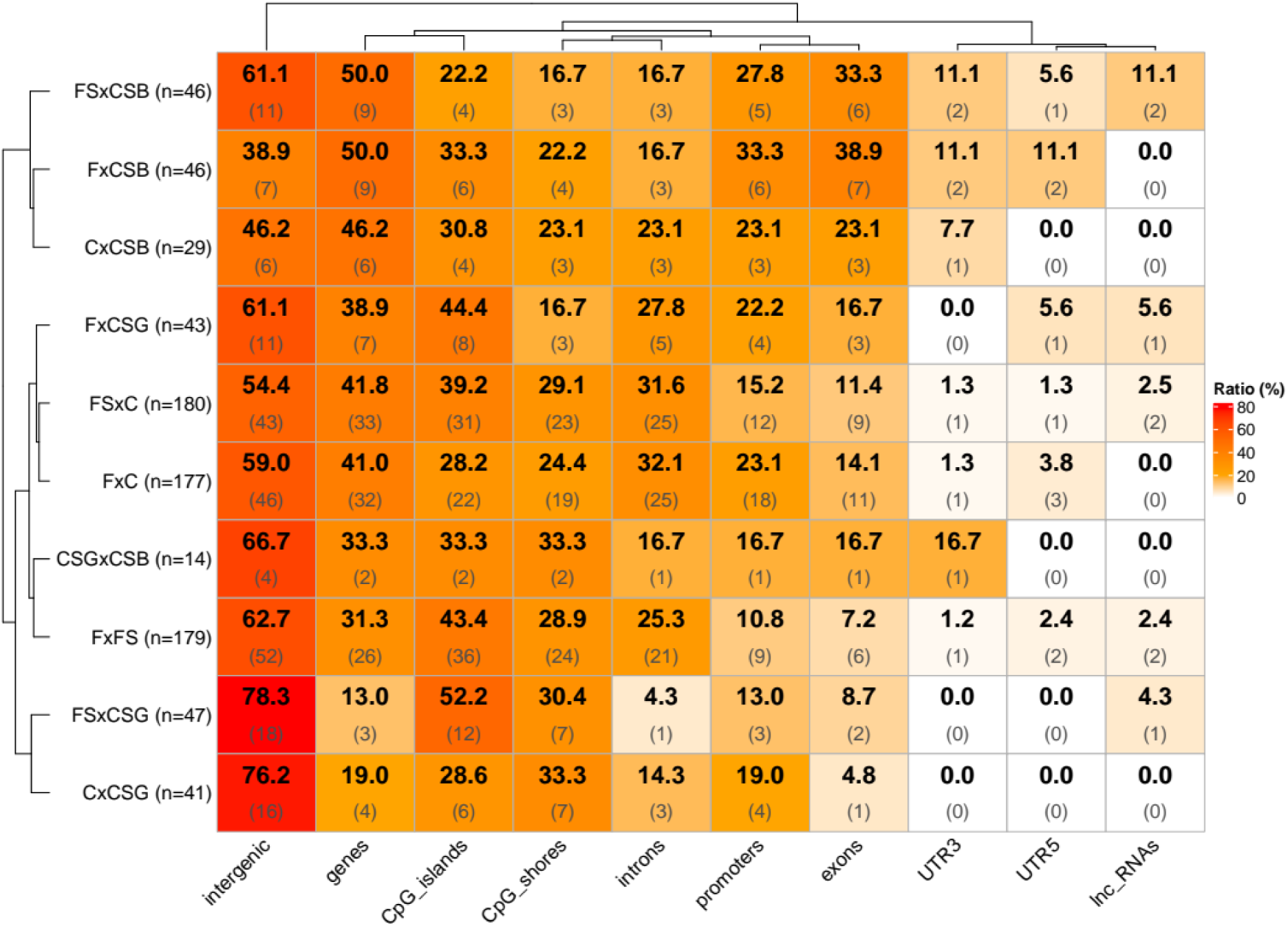
Distribution of relative and absolute DMR counts in known genomic features of the *A. ruthenus* spermatozoa genome for all comparisons. Each cell shows the relative DMR number ratio (the larger black number in bold in the top row) and absolute DMR counts per genomic feature (the smaller grey number in brackets in the bottom row). Clustering was applied both row-wise and column-wise. The Y-axis labels show the comparison names, followed by the total absolute counts of the identified DMRs for each comparison in brackets.

The majority of the DMRs were identified in intergenic regions in all comparisons. DMRs associated with genes were generally predominant in introns rather than exons. The lowest DMR ratio within genes was distributed in UTRs. A large ratio of DMRs is also present within promoters.

Genes associated with significant DMRs (DMGs) were linked to the *A. ruthenus* genome annotation and classified to methylation acquisition (hypermethylated DMRs) or loss (hypomethylated DMRs) for each comparison (see Supplementary Table S5).

The gene lists were also searched for shared genes across all comparisons to reveal common trends in methylome changes and the effect of cryopreservation or separation. Fig. 11 shows the number of shared DMGs among different comparisons for both hypomethylated and hypermethylated regions.

**Fig. 11.**
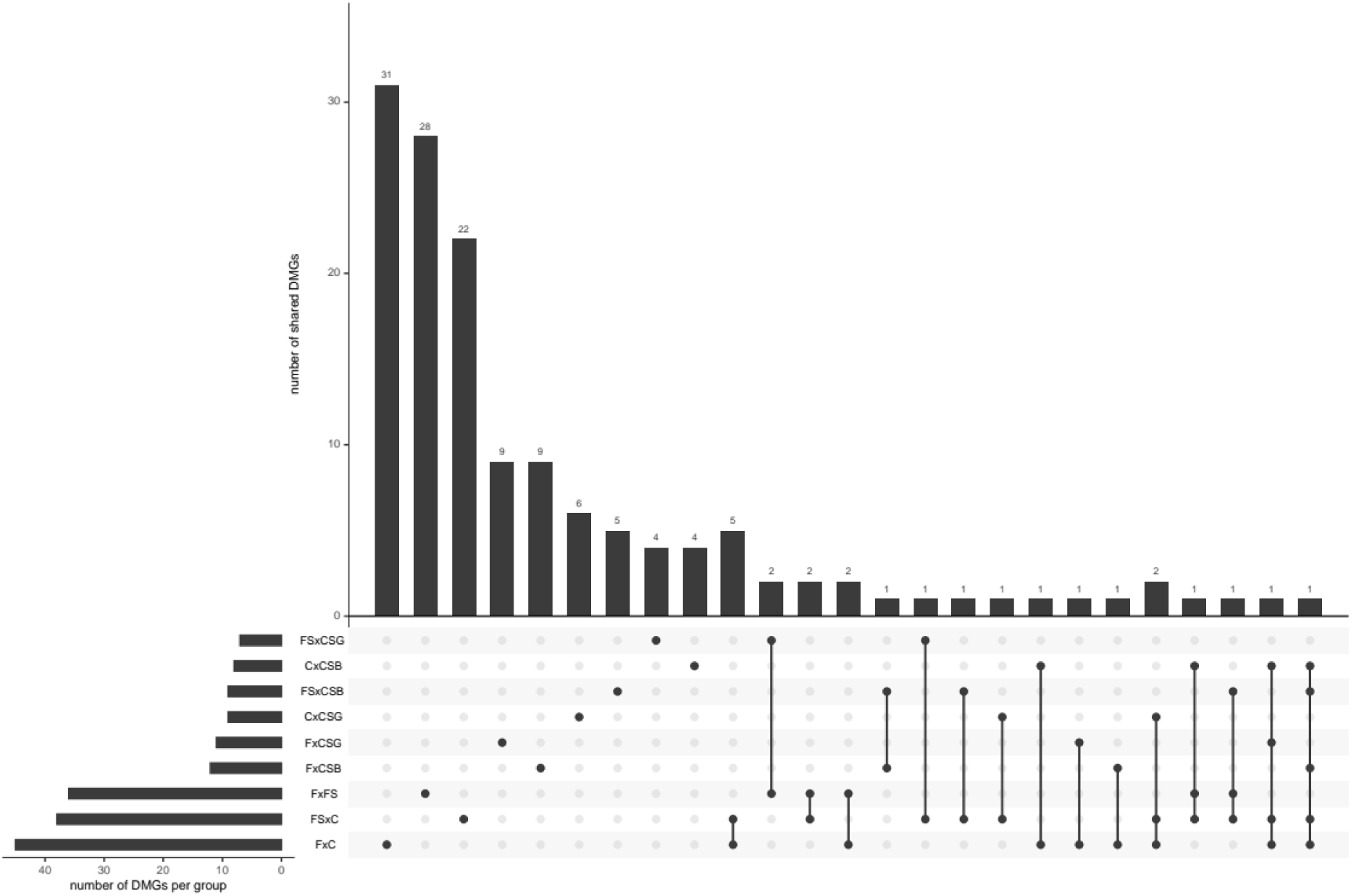
Number of genes shared by one or more comparisons. Each group contains the list of joined hypermethylated and hypomethylated genes. The total number of shared DMGs per comparison is listed above each column. Groups sharing the respective DMGs are indicated in bold dots and connected with a vertical line in the plot under the histogram. Group identity labels are on the left, along with the number of DMGs per individual comparison under the label’s horizontal histogram.

Most DMGs are shared among FxC and FSxC, followed by FxFS and FSxC comparisons. This observation, in line with PCA analysis, shows that the majority of DNA methylation variation is introduced during cryopreservation, and post-thawing phenotype separation does represent a significant source of methylation variability.

The list of shared GeneIDs, corresponding annotations and comparisons sharing the respective GeneIDs is presented in Supplementary Table S6. Most comparisons shared Gene IDs annotated as “uncharacterised genes” or genes associated with basic cellular processes, such as “5S ribosomal RNA” or “transfer RNA glutamic acid (anticodon UUC)”. In the same line, the GSEA did not find any functional enrichment in the list of DMGs in the spermatozoa experimental group comparisons.

#### Experiment 2

##### Bisulfite sequencing of sterlet embryo DNA

Five sterlet spermatozoa experimental groups, Fresh Good (FG), Fresh Bad (FB), Cryopreserved Good (CG), Cryopreserved Bad (CB) and Separated Good (SG) were submitted to whole genome bisulfite sequencing (WGBS), each represented by three biological replicates. The experimental design is outlined in Fig. 2. We received an average of 162 M raw and 120 M clean reads for sperm samples, with an average mapping efficiency of 71.4 %. Details of raw sequencing data quality, raw-read adapter-trimming and quality-filtering, clean read mapping and deduplication statistics are given in Supplementary Tables S7–S9. The all-sample mean read coverage after deduplication is 5.6 x. The graphical representation of the read coverage distribution for each sample is available in Supplementary Fig. S4. The frequency of read coverage distribution for each sample is presented in Supplementary Fig. S5.

The average percentage of methylated cytosines in CpG context (mCpG) in all experimental groups is 70% (Fig. 12), which corresponds to typical vertebrate genome mCpG levels (Elango, Yi, 2008; Tweedie, et al., 1997), although 12% lower compared to mCpG levels in spermatozoa. The all-dinucleotide context cytosine methylation is summarised in Supplementary Table S10.

**Fig. 12.**
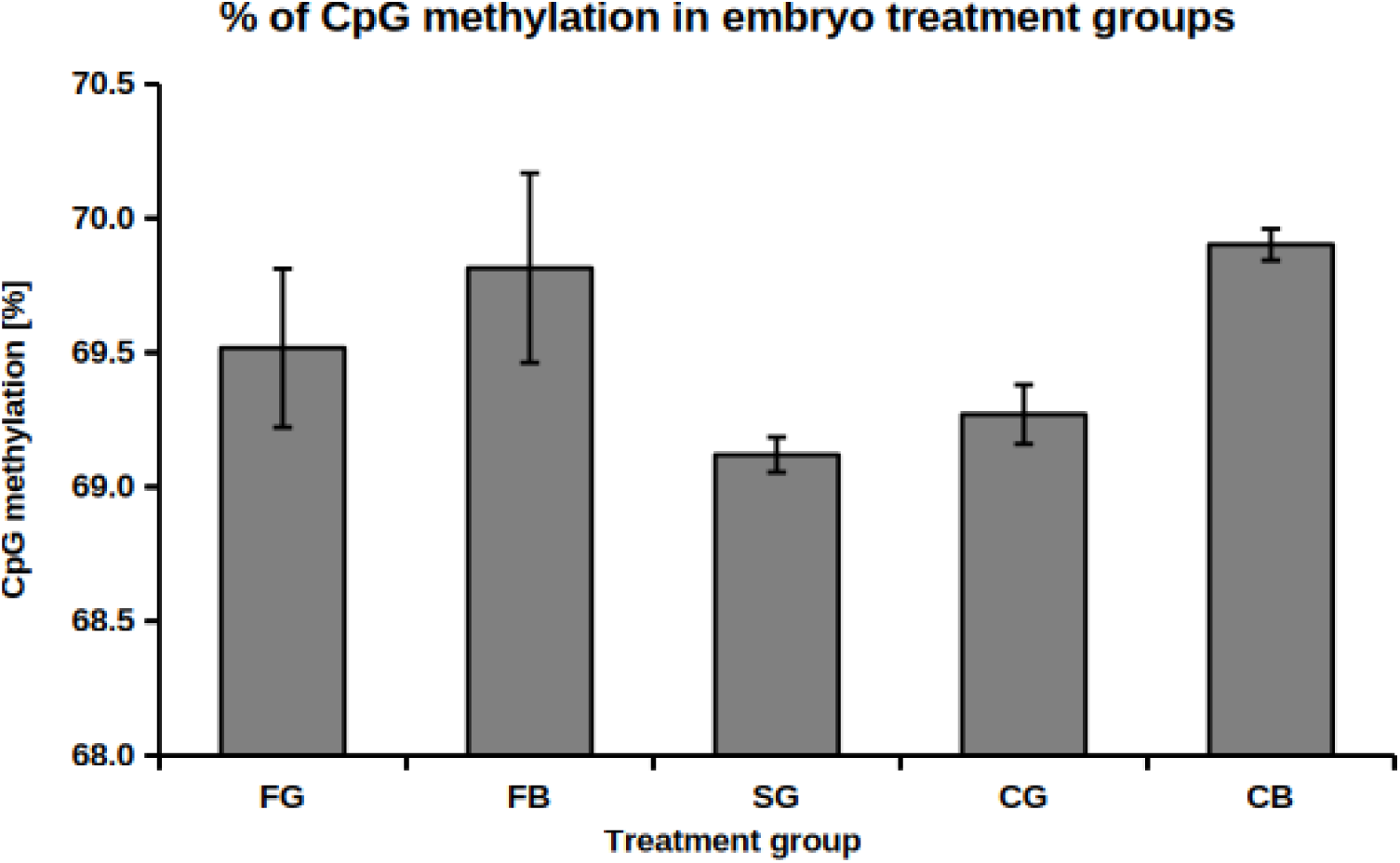
Percentage of CpG methylation in each treatment group, as summarised by the Bismark mapping report. Each value is represented by three biological replications. Treatment groups: FG – Fresh Good, FB – Fresh Bad, SG – Separated Good, CG – Cryopreserved Good, CB – Cryopreserved Bad.

The variance in CpG methylomes among experimental groups was also evaluated with 3D PCA in the same fashion as in the spermatozoa experiment described above (Fig. 13).

**Fig. 13.**
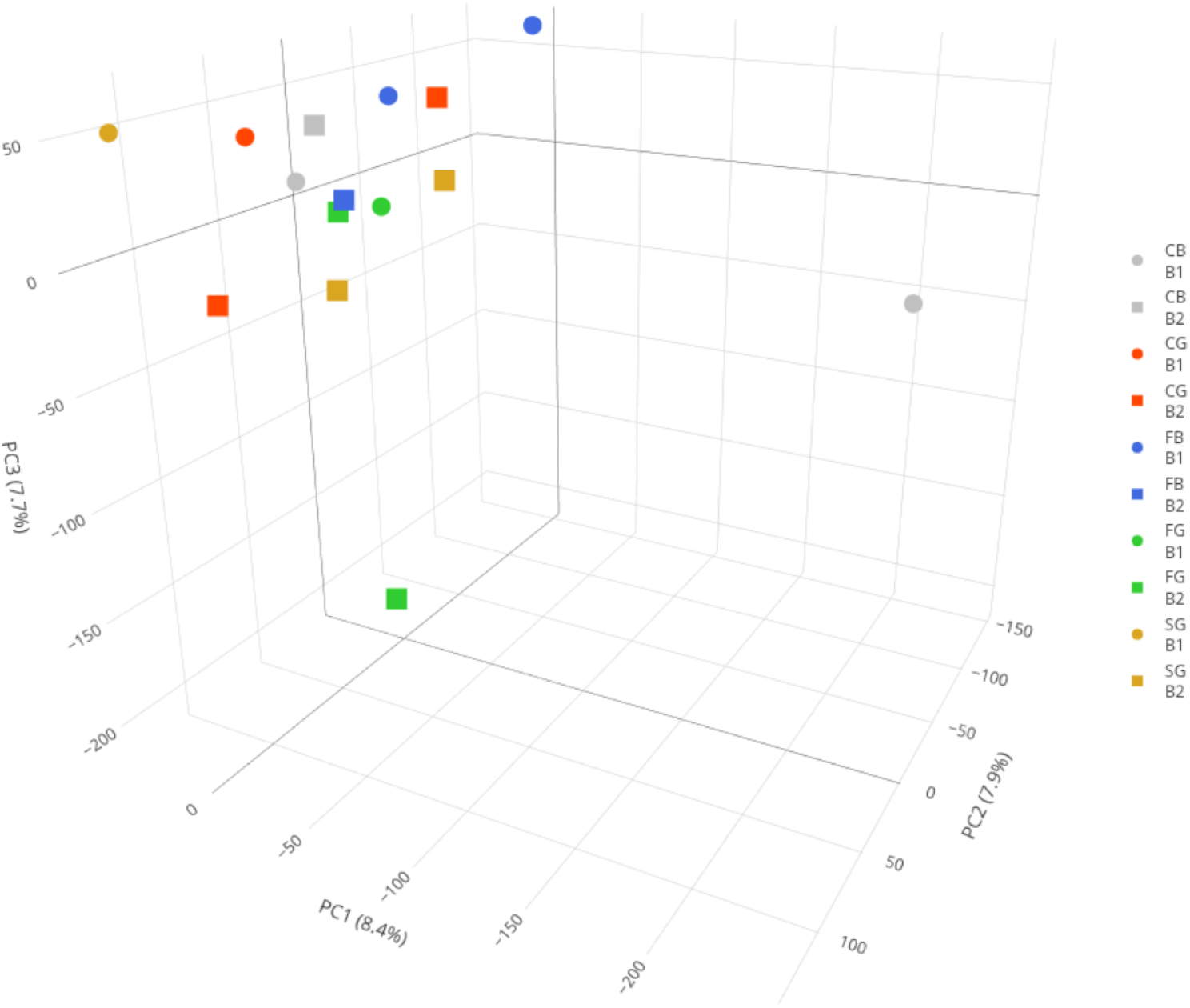
PCA of mCpG counts plotted along 3 major principal components in a 3D scatter plot. The mCpG count matrix of the embryo experimental group was between-sample median-normalised before PCA calculation and plotting. Data points are sample-wise colour-coded and batch effect affiliation is discriminated by data point shape; circle – B1 batch group, rectangle – B2 batch group. Axis labels show % of data variance explained by each PC dimension.

PCA in the 3D space of principal components did not produce any discrete sample clusters. Instead, the majority of data points cluster near zero variance along all 3 axes and show little to no methylome variation among embryo experimental groups.

##### Identification and quantification of differentially methylated regions

DMRs in the CpG context were identified in a pair-wise comparison of seven experimental group pairs. The methylation per cent change and corresponding p-value for each comparison, along with the total number of DMRs and their association with genes and intergenic regions, are presented in Table 2.

**Table 2.** Per cent change of differentially methylated regions (DMRs) for each pairwise comparison as calculated by Defiant*.

| control group | treatment group | percent change [%] | p-value | significance level | tot DMRs | DMRs genes | DMRs intergenic |
| --- | --- | --- | --- | --- | --- | --- | --- |
| FG | FB | 0.2736 | 0.2712 |  | 2 | 0 | 2 |
| FG | CG | -0.5358 | 0.0730 |  | 0 | 0 | 0 |
| FG | CB | 0.3646 | 0.1063 |  | 5 | 1 | 4 |
| FG | SG | -0.4157 | 0.0823 |  | 3 | 0 | 3 |
| SG | CG | 0.1536 | 0.0775 |  | 2 | 0 | 2 |
| SG | CB | 0.7803 | 5.35E-05 | **** | 0 | 0 | 0 |
| CG | CB | 0.6267 | 0.0017 | ** | 0 | 0 | 0 |

\* Each comparison represented by three biological replications is supported by a corresponding p-value and significance level (\* $p < 0.05$ , \*\* $p < 0.01$ ). Absolute numbers of total DMRs, and DMRs associated separately with genes and intergenic regions, are also listed. Treatment groups: FG – Fresh Good, FB – Fresh Bad, CG – Cryopreserved Good, CB – Cryopreserved Bad, SG – Separated Good.

Little to no CpG methylome changes were observed between the studied experimental groups. The global methylome was significantly increased in SGxCB and CGxCB comparisons. However, involved mCpGs were not associated with any specific genomic regions, and thus no DMRs were identified in these and FGxFB comparisons. The absence of significant methylome changes in all comparisons indicates that offspring of spermatozoa exposed to cryopreservation or separation of different physiological forms remain unaffected.

##### Annotation of DMRs

Significant DMRs were mapped to known genomic features in the *A. ruthenus* genome following the spermatozoa annotation pipeline described above. Fig. 14 presents a clustered heat map showing both relative and absolute DMR counts for each comparison and genomic feature. Relative DMR ratios include DMRs shared by overlapping genomic features, and thus the row-wise ratios sum up above 100% per each comparison. Comparisons with zero DMRs were not plotted.

**Fig. 14.**
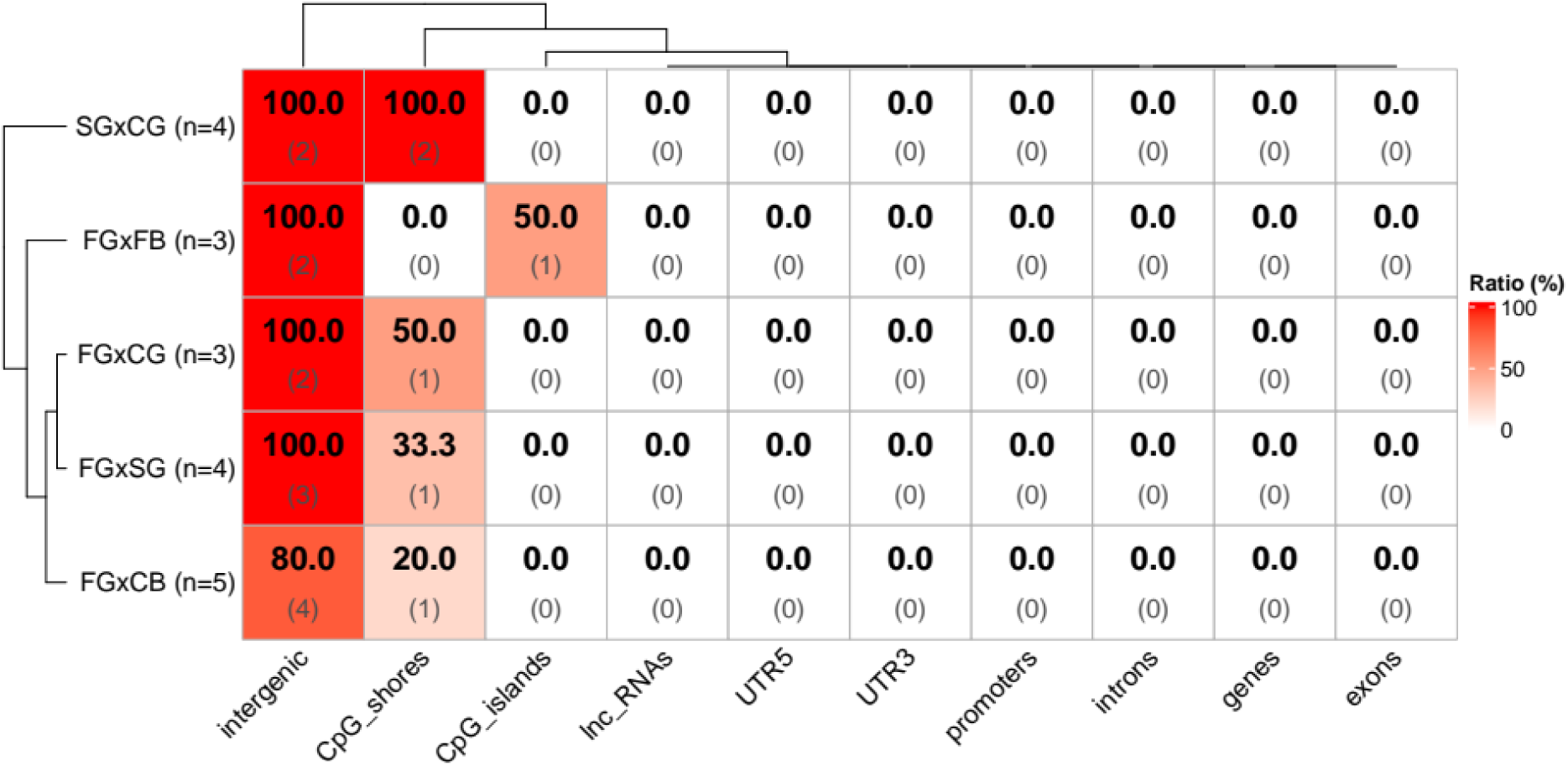
Distribution of relative and absolute DMR counts in known genomic features of the *A. ruthenus* genome for all comparisons. Each cell shows relative DMR ratios (the larger black number in bold in the top row) and absolute DMR counts per genomic feature (the smaller grey number in round brackets in the bottom row). Clustering was applied both row-wise and column-wise. The Y-axis labels show comparisons followed by total absolute counts of identified DMRs per comparison in brackets.

All significant DMRs were found in intergenic regions in all comparisons. More DMRs were mapped to CpG shores than to CpG islands. This correlates with findings that present CpG shores as more dynamic than CpG islands in terms of CpG methylation (Irizarry, et al., 2009; Ziller, et al., 2013).

Defiant identified only two significant DMGs, hypermethylated 117407705 (E3 ubiquitin-protein ligase TRIM35-like) in FGxCG comparison and hypermethylated 117395595 (atrial natriuretic peptide receptor 1-like) in FGxCB comparison.

## Discussion

### Sperm motility

Cryopreservation exposes spermatozoa to multiple stresses, including osmotic imbalance, membrane phase transitions, and oxidative damage, often resulting in reduced post-thaw motility and fertilisation success (Fuller, et al., 2004; Holt, 2023). These effects are well documented in cryobiology and fish sperm studies, generally indicating deterioration of sperm performance after freezing (Asturiano, et al., 2017; Tiersch, Green, 2011). However, in sturgeons, reported outcomes vary widely depending on species, protocol, and initial sperm quality, with motility parameters showing decreases (Dzyuba, et al., 2012; Horváth, et al., 2008b; Sieczyński, et al., 2015), no change (Dzyuba, et al., 2010; Judycka, et al., 2015), or even increases (Boryshpolets, et al., 2011) compared to fresh controls.

In this study, cryopreservation caused a marked and consistent reduction in motility (MOT) in non-separated sperm, whereas Percoll density-gradient centrifugation (DGC) significantly improved MOT both before and after freezing. This enrichment of a high-motility subpopulation aligns with previous findings in sturgeon, where motile fractions retained high viability and minimal proteomic changes relative to fresh sperm (Horokhovatskyi, et al., 2018). CASA analysis revealed strong treatment effects on MOT, while most kinematic parameters (VCL, VAP, VSL, LIN, WOB, BCF) showed only sporadic differences, suggesting that Percoll primarily selects sperm capable of activation and sustained movement rather than those with distinct kinematic traits. Cluster analysis supported this interpretation, showing similar temporal dynamics across groups.

Importantly, motility was assessed during the time window relevant for in vitro fertilisation, ensuring biological relevance. Differences from Horokhovatskyi et al. (Horokhovatskyi, et al., 2018), where Percoll also enriched sperm with higher kinematic values, likely reflect male variability in cryoresistance (Kopeika, Kopeika, 2008). Unlike patterns reported in non-sturgeon species, where cryopreservation reduces the abundance of sperm fractions with high velocity (Bravo, et al., 2020; Parks, Graham, 1992; Pérez-Atehortúa, et al., 2021), our findings show that Percoll DGC increased motility in sterlet, consistent with results in other fish such as catfish (Rhamdia quelen) (Pérez-Atehortúa, et al., 2021) and Atlantic salmon (Salmo salar) (Bravo, et al., 2020). Overall, Percoll DGC enhances functional sperm quality by removing non-activating cells rather than altering motile sperm kinematics, providing a robust basis for subsequent analyses of the reproductive potential and DNA state in motile, presumably non-cryodamaged sperm.

### Reproductive performances

The influence of sperm cryopreservation on progeny performance in fishes remains controversial, with reported outcomes ranging from negative to neutral or even positive effects on offspring development. Notably, in several species, fertilisation and hatching rates obtained using cryopreserved sperm are often comparable to those achieved with fresh sperm, despite marked reductions in post-thaw sperm motility and membrane integrity (Figueroa, et al., 2016; Horváth, et al., 2003; Linhart, et al., 2000). This apparent resilience of early developmental endpoints indicates that fertilisation success alone may an insufficiently sensitive indicator of cryopreservation-induced sperm damage.

Evidence from cyprinids and salmonids indicates that cryopreservation can influence progeny performance in divergent ways at later developmental stages. In salmonids such as Atlantic salmon and brown trout, the use of cryopreserved sperm has been linked with reduced larval growth or altered performance, with outcomes depending on the cryoprotectant and freezing protocol applied, even when fertilisation rates remain unaffected (Dziewulska, et al., 2011; Nusbaumer, et al., 2019). In contrast, study in common carp has shown that growth performance during pre-nursing and grow-out periods phases can be comparable between progeny produced using fresh and cryopreserved sperm (Bokor, et al., 2024). These contrasting findings demonstrate that cryopreservation-induced effects on progeny are not universal but instead appear to be species-specific and shaped by the interaction between the type and extent of sperm damage, the developmental stage examined, and the environmental and husbandry conditions under which offspring are reared.

Mechanistically, cryopreservation is known to induce oxidative stress, membrane lipid rearrangements, and metabolic disturbances in spermatozoa (Alvarez, Storey, 1992), which may compromise DNA integrity or epigenetic stability without preventing fertilisation (Cabrita, et al., 2010). Such sublethal damage may allow functionally compromised spermatozoa to contribute to fertilisation and subsequently influence embryonic or larval development. Increasing evidence further suggests that sperm populations respond heterogeneously to cryopreservation, with subsets of spermatozoa retaining higher functional competence after freezing and thawing (Gallego, et al., 2017). This heterogeneity provides a plausible explanation for the frequent discrepancy between fertilisation success and later progeny performance reported across studies.

Together, our findings and existing literature supports the view that sperm cryopreservation in fishes can influence progeny performance through subtle, population-level effects that are not captured by fertilisation or hatching rates alone. These findings underscore the importance of considering sperm heterogeneity and post-thaw sperm selection when evaluating cryopreservation protocols, particularly in aquaculture and conservation contexts where progeny quality is a critical outcome.

Given these variable and often subtle effects of cryopreservation on progeny performance, an important question is whether such functional differences are accompanied by underlying molecular changes in the sperm epigenome. Because spermatozoa carry regulatory information in addition to DNA sequence, cryo-induced stress has the potential to alter epigenetic marks that could, in principle, influence early development even when fertilisation outcomes appear normal. Assessing whether cryopreservation and post-thaw sperm selection modify sperm DNA methylation—and whether any such changes persist in embryos—therefore provides a critical mechanistic link between observed sperm heterogeneity and potential downstream effects. Our study directly addresses this by evaluating methylation patterns in sterlet sperm and their transmission to early embryos.

### DNA methylation

DNA methylation is a mechanism which evolved to facilitate rapid adaptation to immediate external stimuli of the host organism, regulating appropriate cellular pathways at the level of gene expression (Feil, Fraga, 2012; Flores, et al., 2013; Law, Holland, 2019). Cryopreservation, in this respect, represents an intense external stimulus and, at the same time, a significant source of stress.

In this study, we describe in-captive breeding techniques in A. ruthenus aquaculture and address their effect on the physiological parameters and methylomes of sterlet spermatozoa and their progeny.

Global DNA methylation varies from 72 to 98 % among different fish species under physiological conditions (Chen, et al., 2021; Feng, et al., 2010; Freij, et al., 2024; He, et al., 2022; Hou, et al., 2023; Pan, et al., 2021; Wellband, et al., 2021; Zhou, et al., 2019). Previous studies investigating methylation in other Acipenser species have shown markedly variable levels of global genome methylation ranging from 16 to 78 % across different tissues and physiological conditions (Earhart, et al., 2023; Whitaker, et al., 2018).

Methylome analysis of the sterlet genome in our work revealed 82 and 71.4 % of mCpGs in spermatozoa and embryo genomic DNA, respectively. This observation is consistent with global DNA methylation levels in both sperm and embryo reported in carp (80 % in spermatozoa and oocyte) (Cheng, et al., 2021; Cheng, et al., 2024), zebrafish (95 % spermatozoa, 75 % oocyte) and medaka (91 % spermatozoa, 80 % oocyte) (Jiang, et al., 2013; Potok, et al., 2013; Wang, Bhandari, 2019). Observed sperm genome hypermethylation aligns with a well-known phenomenon of high genome condensation to ensure its stability during fertilisation (Güneş, Kulaç, 2013).

The dimensionality reduction and projection of the mCpG count covariance matrix in the 3D PCA space discriminated three distinct data-point clusters associated with Fresh, Fresh separated, and Cryopreserved groups. This clustering pattern confirms both biological replication consistency and a linkage between cryopreservation and/or phenotype separation with spermatozoa DNA methylation pattern. The inhomogeneous data point cloud formed by the CSG and CSB group samples implies that methylation does not induce distinct phenotypes in the post-cryopreserved spermatozoa population.

Formation of spatially remote treatment/phenotype-specific group clusters also documents that cryopreservation and phenotype separation represent distinct physiological conditions that drive and/or are induced by specific changes in DNA methylation. Here, identifying causality in the observed pattern is also crucial. While cryopreservation represents an environmental stimulus that alters host cell DNA methylation, different physiological phenotypes can arise in response to altered DNA methylation changes. Our data suggest that this is also the case in the sterlet spermatozoa experimental groups.

Separation of physiological phenotypes after cryopreservation was not associated with methylome variability, as evidenced by the diffuse distribution of CSG and CSB data points within the 3D PCA space. The data distribution and cluster formation are explained by 29.5 % of the total mCpG count variance as the sum of all three PC dimensions. We had to project methylation variability into three PC dimensions, as two dimensions did not show enough variability for efficient sample cluster definition.

The PCA was also used to detect potential batch effects on the methylome variance among experimental groups. As the groups, which may have arisen from potential batch effect-induced variability, are randomly distributed across sample groups, we can conclude that methylome variability forming sample-group clusters was not driven by batch effect but only by sample treatment.

We performed DMR analysis using pairwise comparisons of 10 relevant experimental group pairs to identify cryopreservation- and phenotype-associated effects in DNA methylation at the level of specific genomic regions and genes. These data, in combination with DMR analysis in the embryo, should determine whether methylation changes observed in spermatozoa are translated into quality and physiology in their progeny.

DMR analysis in spermatozoa identified small but significant changes in six pairwise comparisons. The highest significant mCpG methylation change was observed between fresh (F group) and cryopreserved spermatozoa in the high-motility fraction (CSG group), followed by FxCSB and FSxCSG group comparisons. Of note, the largest statistically significant methylome changes were observed in high- and low-motility fraction experimental groups compared with cryopreserved experimental groups. It seems that cryopreservation does not influence the methylome, as demonstrated by the insignificant methylome changes in FxC versus significant in FxCSG, FxCSB comparisons, and, analogically, insignificant in FSxC versus DMGs identified in each comparison were not overrepresented in any GO category by GSEA. Along the same lines, a search for DMG intersections did not cluster comparisons by cryopreservation or separation into motile and non-motile fractions. It is interesting to note that a high proportion of DMGs were shared across multiple comparisons relative to the total number of identified DMGs, suggesting that these genes may be more prone to DMGs in spermatozoa during the artificial fertilisation process. For example, differentially methylated Histone H3 is often associated with stress response in several studies (Delaney, et al., 2018; Nagagaki, et al., 2024; Nunez-Vazquez, et al., 2022; Nunez-Vazquez, et al., 2025). Its differential methylation in transcriptionally inactive sperm can alter the chromatin architecture of the spermatozoa genome, influence accessibility of affected genomic regions and influence gene expression in offspring.

DMRs in the remaining four comparisons were not statistically significant, and their origin can be attributed to spontaneous changes or random internal or external stimuli which naturally occur in fish spermatozoa.

DMRs for both the spermatozoa and embryo experimental groups were linked to the coordinates of the most relevant known genomic features. The majority of DMRs were located in intergenic regions of the sterlet spermatozoa genome, followed by genes, with introns being preferentially occupied over exons in most comparisons.

DNA methylation mechanism in eukaryotic cells was adopted from prokaryotes and adapted for repetitive DNA silencing and later for gene expression regulation in complex eukaryotic genomes (Ehrlich, Lacey, 2013; Jjingo, et al., 2012). Permanently methylated intergenic regions are thus often associated with mobile elements and repetitive DNA sequences to repress their harmful spread throughout the genome. Intergenic regions are also niches for regulatory sequences, acting as distant controls of gene expression, such as transcription enhancers, silencers, or insulators, and their differential methylation also plays an important role in gene expression regulation. Many studies found incorrect methylation states to cause aberrant gene expression leading to physiological malformations and diseases (Fan, et al., 2024; Jeziorska, et al., 2017; Shenker, Flanagan, 2012; Yan, et al., 2016).

The presence of methylated CpGs in genes, particularly in gene bodies, is a typically a transcriptionally independent feature of vertebrate genomes (Jjingo, et al., 2012). In contrast, CpGs in promoters are organised in CpG islands, and their methylation correlates with transcriptional repression (Chodavarapu, et al., 2010; Lister, et al., 2013; Shukla, et al., 2011). Promoter methylation changes correlate with altered gene expression in the affected experimental groups. The highest incidence of DMRs within promoter regions relative to other genomic features was identified in FSxCSB and FxCSB comparisons. The highest absolute counts were observed in FxC and FSxC comparisons. This implies that cryopreservation induced the highest DMR numbers in gene promoters, potentially altering the expression of associated genes. It is a matter of debate whether these relatively low numbers of DMRs could significantly affect gene regulation. Such hypothesis have to be investigated in more details in relation to CpG colocalization with specific sequence motives, their spatial distribution or their combination with other marks associated with transcription regulation within promoters and also with the fact that mCpGs sequence and spatial contexts play more important role in gene expression regulation than their density (Sapozhnikov, Szyf, 2021; Taryma-Leśniak, et al., 2024).

DMR annotation revealed a discordance between the Defiant and the Genomation tool results. While Defiant identified 2 DMGs in 2 comparisons, Genomation did not find any DMRs overlapping with genes in the *A. ruthenus* genome. This kind of disagreement can arise from how the gene coordinates are defined. For example, some tools include the promoter region in the gene definition, while others do not. Also, promoter coordinates can be defined differently. Some tools may consider only protein-coding genes. There are several sources of genome annotations, and both tools may use different sources even when using the same genome assembly and version. In this case, Defiant by default defines gene promoters as regions 10,000 nucleotides upstream of gene coordinates, extracted from the associated genome annotation file for DMR analysis following more generous promoter coordinates definition as in studies focusing specifically on gene expression where authors may want to extend this definition to include all regions that could be involved in gene expression regulatory networks (Kaplun, et al., 2016). Contrarily, *A. ruthenus* genome annotation performed in our study used generally recommended and more widely used promoter coordinate spans of ∼500-1000 bp upstream of the TSS (FitzGerald, et al., 2004; Tabach, et al., 2007; Zhang, 2007).

DMGs identified in each comparison were not overrepresented in any GO category by GSEA. Along the same lines, a search for DMG intersections did not cluster comparisons by cryopreservation or separation into motile and non-motile fractions.

Many cellular responses are triggered by cold stress, and the immune system is among the first to be alerted (Abram, et al., 2017). Interleukin 12B (IL12B), an integral part of pro-inflammatory cytokine-mediated immune response, was found to be differentially expressed in response to cold stress in studied vertebrates (Hangalapura, et al., 2006; Monroy, et al., 1999). Our study found Interleukin 12B (IL12B) on the list of differentially methylated genes in cryopreserved spermatozoa, and its altered methylation state can affect cold-temperature sensitivity and progeny adaptation when inherited.

Several genes associated with lipid metabolism were also differentially methylated. Lipid metabolism pathways are typically induced by cold stress, as in the case of cAMP-specific phosphodiesterase 4D (PDE4D), which activates brown adipose tissue and lipolysis, leading to thermogenesis, or Butyrophilin subfamily 1 member A1 (BTN1A1-like), which plays a role in lipid metabolism, mainly lipid secretion, and the immune response. BTN1A1-like is associated with membrane lipid composition and can affect spermatozoa membrane fluidity and integrity during spermatogenesis in progeny. Although neither of two genes has been directly linked to cold-induced stress, their differential expression was associated with more general stress responses in vertebrates (Dou, et al., 2020; Hu, et al., 2016; Malinowska, et al., 2017), and their cold-stress association is to be revealed by more detailed studies focused on temperature-induced stress.

Phosphoenolpyruvate carboxykinase (PCK1, cytosolic [GTP]) (PCK1) and PDE4D are enzymes involved in energy regulation, motility activation, and membrane fluidity, important for sperm motility. Differential methylation in these genes can affect sperm maturation in fish gonads and alter spermatozoa motility and cold sensitivity in the adult male progeny.

Specifically, PCK1 expression was altered in metabolic adaptive response to low temperatures in several fish species (Guo, et al., 2025; Miao, et al., 2021; Windisch, et al., 2011). TNFRSF10B-like and HES-5-like (Notch signalling) are recognized as general markers of active cellular stress response (Li, et al., 2015; Sullivan, et al., 2020; Zhao, et al., 2025).

The above observations imply that spermatozoa responded to cryopreservation by differential methylation of genes associated with both general stress pathways and low-temperature-mediated cellular stress responses. Although these genes were significantly differentially methylated in our data, none were enriched in any GO category, and their overall effect on potential changes in cellular response must be considered.

Due to transcriptional silencing of the spermatozoa genome, differential methylation in spermatozoa does not directly influence spermatozoa fitness. Instead, differential gene methylation in spermatozoa may influence DNA stability, chromatin organisation or histone retention, all of which shape sperm nuclear architecture and can potentially influence embryo gene activation upon fertilisation.

Regarding changes in embryo fitness and methylation patterns, we did not observe any significant methylation changes in the embryo experimental groups. In contrast to spermatozoa, the methylation changes were not observed at either the whole methylome level or in specific genomic regions in the embryo.

As a regulator of gene expression, methylation also functions in a tissue-specific manner, resulting in tissue-specific methylation patterns (Lokk, et al., 2014; Maunakea, et al., 2010; Wiench, et al., 2011; Xie, et al., 2013; Zhang, et al., 2013; Zhou, et al., 2017) and embryonic tissue is not an exception (Isagawa, et al., 2011; Loyfer, et al., 2023; Sørensen, et al., 2010). The mixture of tissue-specific methylation patterns within individual embryos could explain the absence of a specific methylation pattern among experimental groups of sterlet embryos. This hypothesis should be verified by high-resolution analyses using advanced techniques such as single-cell methylation analysis (Farlik, et al., 2015; Liu, et al., 2023; Nichols, et al., 2025; Nichols, et al., 2022), an approach only recently implemented in zebrafish (Burgos-Ruiz, et al., 2026).

The extent of DNA methylation heritability also differs among organisms. Unlike the well-studied methylation inheritance in mammals, little is known about methylation dynamics and its inheritance in fish. The developmental dynamics of DNA methylation was described during embryogenesis in a few fish species (Jiang, et al., 2013; Potok, et al., 2013; Wang, Bhandari, 2019). Although the extent of methylation clearance in fish embryos is significant, it does not reach that of mammals, undergoing almost complete parental methylome reset (Morgan, et al., 2005), providing space for certain transgenerational epigenetic inheritance in fish (Matlosz, et al., 2024; Ortega-Recalde, et al., 2019; Ross, et al., 2023).

DNA methylation is a highly dynamic epigenetic mark that confers phenotypic plasticity and the ability to respond to immediate environmental changes on its hosts. Rapid and extensive changes in DNA methylomes were recorded in response to different kinds of environmental stimuli in fish (Anastasiadi, et al., 2017; Brionne, et al., 2023; Cavalieri, Spinelli, 2017; Liu, et al., 2025; Ma, et al., 2025; Navarro-Martín, et al., 2011; Sánchez-Baizán, et al., 2025; Strömqvist, et al., 2010; Wang, et al., 2009; Welengane, et al., 2025; Yao, et al., 2022).

Effects of environmentally-induced epigenetic stimuli in fish were found to be transgenerational (Abdelnour, et al., 2024). However, their inheritance can vary by organism and character of environmental stimulus, inducing corresponding methylation changes. For example, rearing temperature affected parental methylome in brook charr but was not inherited by its offspring (Venney, et al., 2022). Contrarily, temperature-induced changes in the methylome were reflected in offspring phenotype in rainbow trout (Brionne, et al., 2023). Methylation changes induced by hydrogen sulfide-rich springs were inherited by progeny of Atlantic molly (Kelley, et al., 2021) and similar xenobiotic-induced methylation changes were also found heritable in zebrafish, a vertebrate model for ecotoxicology research, where a myriad of various environmental pollutants and toxins were investigated (Kamstra, et al., 2015; Terrazas-Salgado, et al., 2022).

Aquaculture management and fertilisation techniques represent a repertoire of external stimuli causing gene expression and/or DNA methylation-induced changes in spermatozoa phenotypes (Makvandi-Nejad, Moghadam, 2023). Many studies reported direct evidence of altered methylation patterns in fish hatcheries compared to wild populations, hence questioning fish conservation and reintroduction projects (Gavery, et al., 2019; Griffiths, et al., 2025; Koch, et al., 2023; Nilsson, et al., 2021; Podgorniak, et al., 2022; Rodriguez Barreto, et al., 2019; Venney, et al., 2025). Same considerations must also be taken into account in fish farming aquaculture management (Zhang, et al., 2023), where the pursuit for optimised breeding protocols, high fish sperm motility and fertilisation capacity and maximised yields of high-quality fish progeny can be hindered by negative effects of animal rearing in captivity and artificial fertilisation techniques.

Many studies have already demonstrated that spermatozoa handling, storage and cryopreservation alter the methylation pattern of the spermatozoa genome, ultimately leading to changes in spermatozoa phenotypes in different fish species (Cheng, et al., 2021; Cheng, et al., 2024; El Kamouh, et al., 2023; Narud, et al., 2023; Pappas, et al., 2025).

In our project, we focus on optimising *A. ruthenus* fertilisation techniques. We aimed to uncover the source of phenotypic variability resulting from the separation of the motile fraction after cryopreservation, at the level of genome regulation.

Our data found cryopreservation and motile fraction separation to be associated with distinct and significant but small changes in methylation pattern in sterlet spermatozoa, which is in line with previous findings in goldfish, zebrafish and rainbow trout (Depincé, et al., 2020; El Kamouh, et al., 2023).

However, this differential methylation pattern was not inherited in progeny, nor did it correlate with embryo physiological phenotypes. On the contrary, a recent study found that paternal-effect genes primed by cryopreservation were manifested in progeny phenotypes (Panda, et al., 2024). Although the study examined this effect on gene expression level and phenotypic traits, the authors highlight DNA methylation as the most likely priming mechanism for transgenerational carryover of the observed paternally induced phenotypes.

Our study presents the first methylation analysis of the *A. ruthenus* genome. The main limitation of our project is low sequencing coverage, with 8.8x and 5.6x mean read coverage for spermatozoa and embryo genomes, respectively. Our DMR analysis used the minimum recommended CpG coverage filter of ≥ 10. This strict filter excluded CpGs which did not receive sufficient reads in our final dataset. Therefore, the lowly covered genomic regions were excluded from the analysis and could not contribute to finding a potential correlation between DNA methylation, cryopreservation and motile spermatozoa fraction separation. Lowering CpG coverage filtering, as in a previous study that also suffers from low genome coverage (Wellband, et al., 2021), would threaten data reliability and could lead to many false-positive observations and bias the entire analysis. Future studies in this field should secure higher genome coverage to receive more informative and well-supported data.

## Conclusions

Density-gradient centrifugation effectively enriched a high-motility sterlet sperm fraction, improving fertilisation success by reducing malformed larvae. Cryopreservation and sperm selection introduced small but detectable DNA methylation changes in spermatozoa, yet these alterations were not reflected in embryo methylomes and showed no association with developmental outcomes. However, the low sequencing depth in our study does not provide sufficient evidence to draw firm conclusions about the potential effects of sperm methylation changes on progeny. This study contributes to ongoing efforts to optimise sterlet breeding techniques and represents the first whole-genome methylome analysis at nucleotide resolution in *A. ruthenus*. Future work with higher sequencing coverage, integrated gene expression analyses, and broader phenotypic assessments is needed to clarify links between cryopreservation, methylation, and offspring quality. To the best of our knowledge, this study is the first to provide insight into the whole-genome methylome at nucleotide resolution in the Acipenseridae family and the first to publish the DNA methylome of the *Acipenser ruthenus* species.

## Acknowledgements

The study was financially supported by the Czech Science Foundation (No. GACR 22-14069S). Part of the work was carried out with the support of VVI CENAKVA Research Infrastructure (ID 90238, MEYS CR, 2023–2026).

Computational resources were supplied by the Ministry of Education, Youth and Sports of the Czech Republic under the Projects CESNET (Project No. LM2015042) and CERIT-Scientific Cloud (Project No. LM2015085) provided within the program Projects of Large Research, Development and Innovations Infrastructures.

