## Supplementary figures and images for "Improved Sperm Quality and Cryo-Induced Epigenetic Changes in Sterlet via Density-Gradient Sorting"

### Supplementary Fig S1

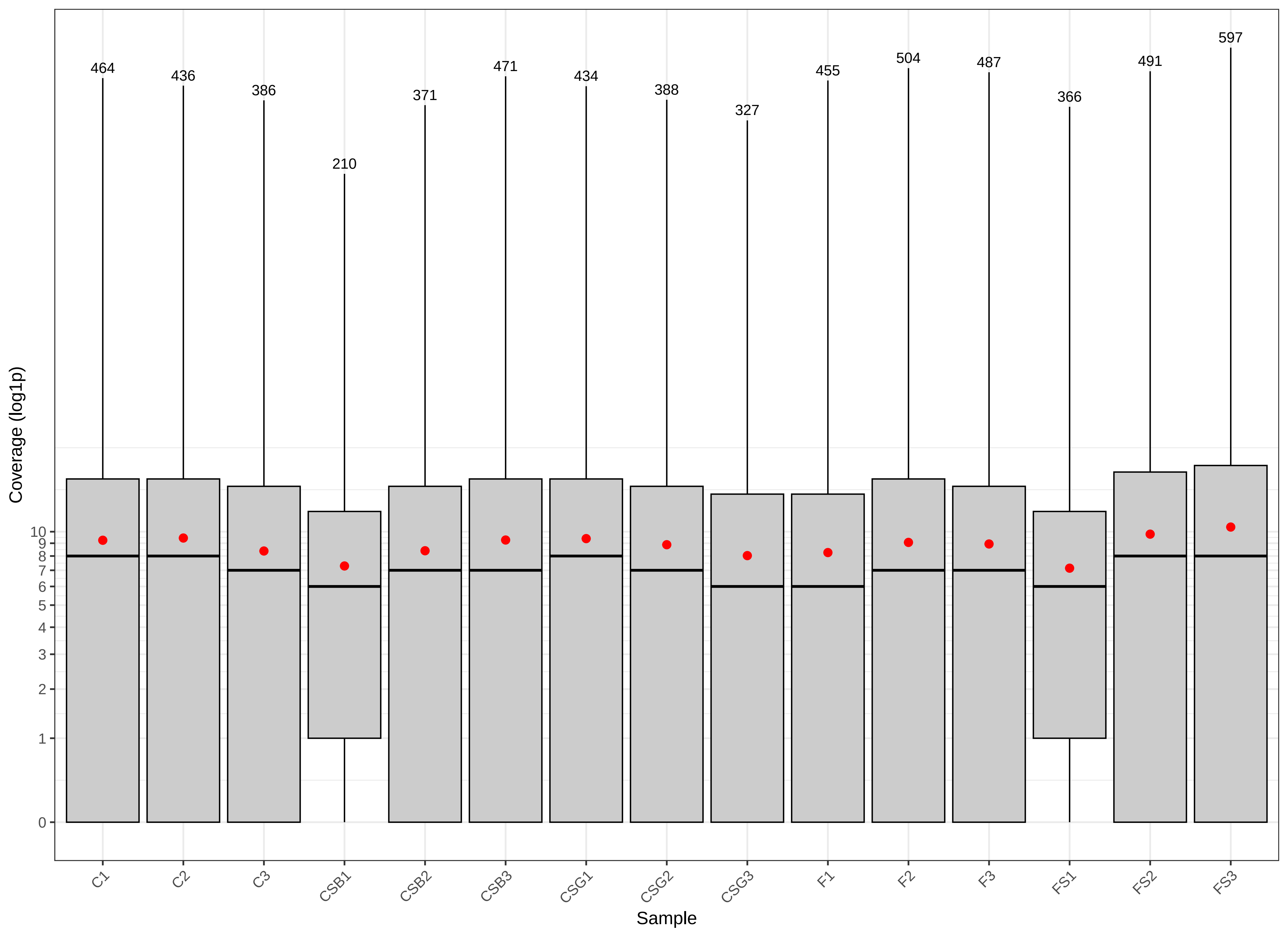

### Supplementary Fig S2

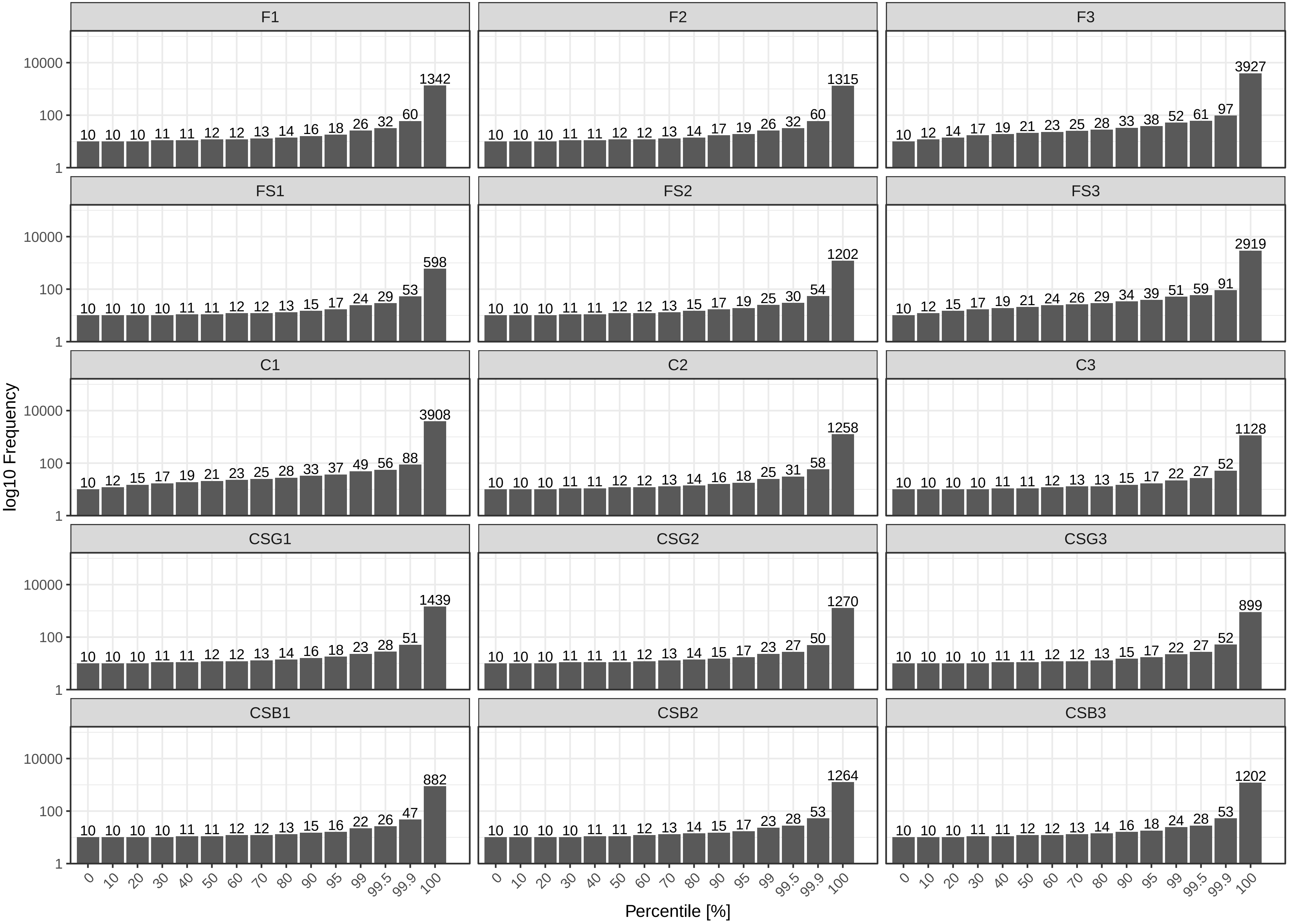

### Supplementary Fig S3

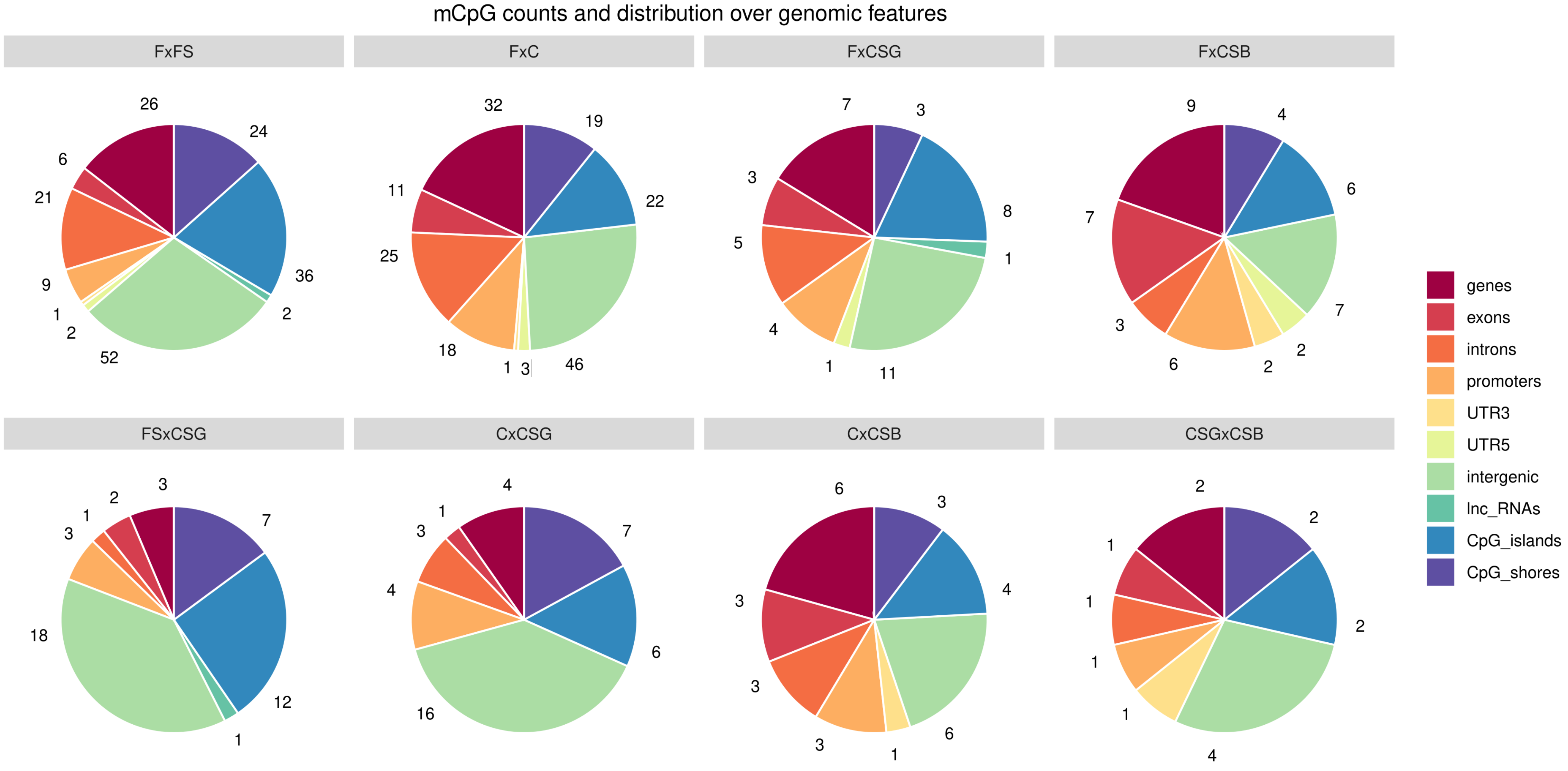

### Supplementary Fig S4

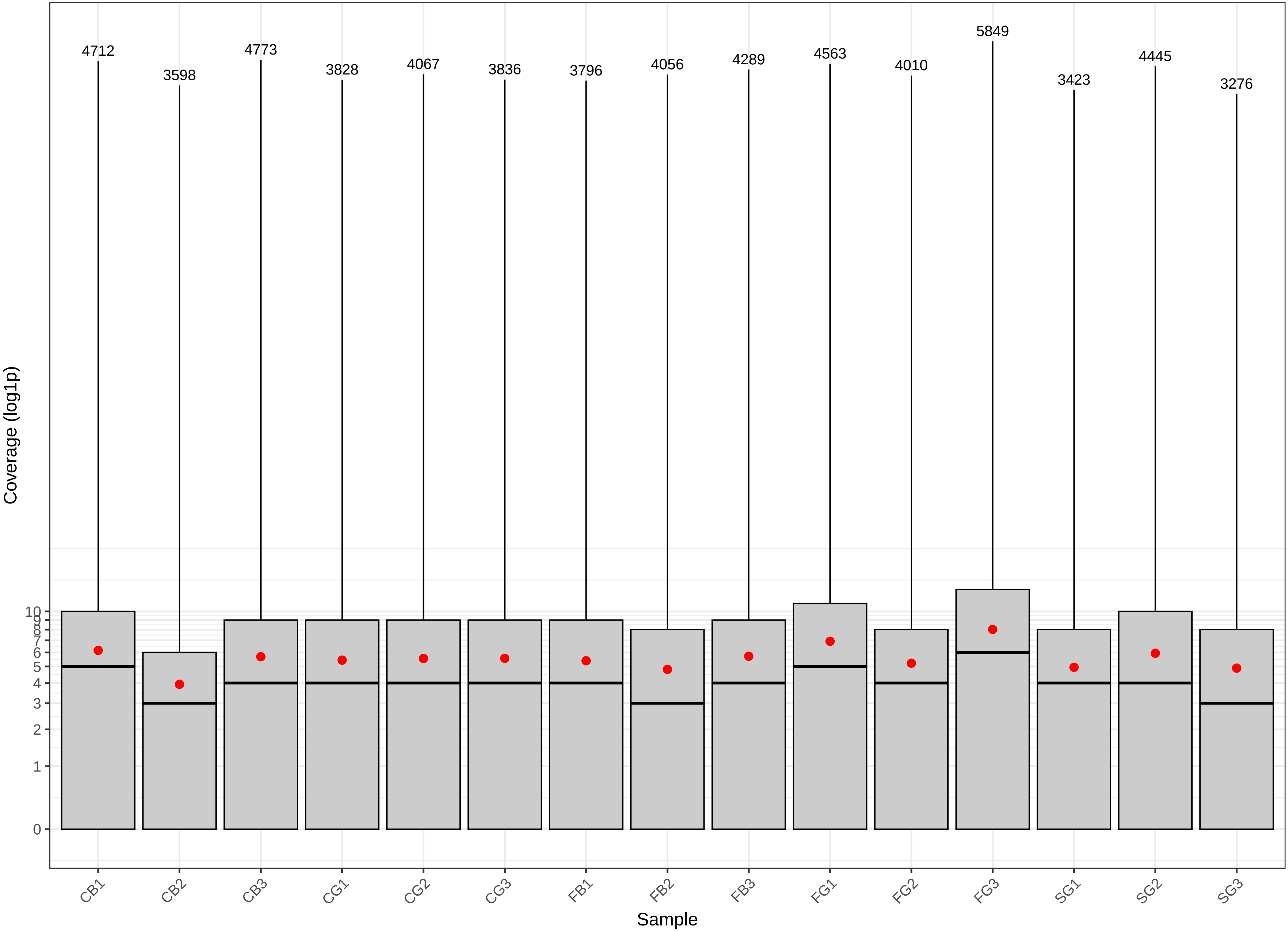

### Supplementary Fig S5

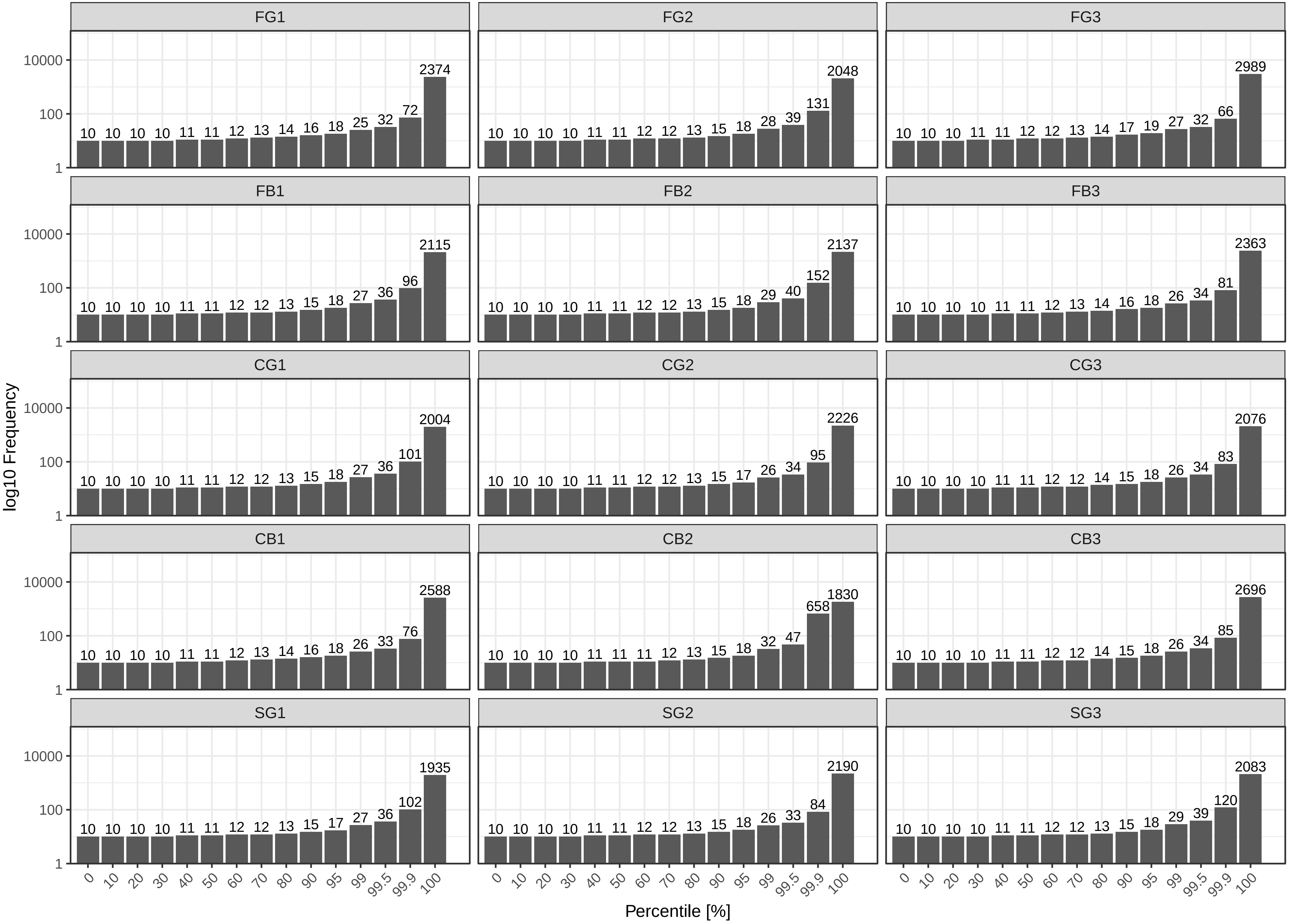
